# Interplay of the folded domain and disordered low-complexity domains along with RNA sequence mediate efficient binding of FUS with RNA

**DOI:** 10.1101/2022.11.09.515892

**Authors:** Sangeetha Balasubramanian, Shovamayee Maharana, Anand Srivastava

## Abstract

RNA binding ability of Fused in Sarcoma (FUS) protein is crucial to its cellular function. Our molecular simulation study on FUS-RNA complex provides atomic resolution insights into the observations from biochemical studies and also illuminate our understanding of molecular driving forces that mediate the structure, stability, and interaction of RRM and RGG domains of FUS with a stem-loop junction RNA. We observe a clear cooperativity and division of labour among the ordered (RRM) and disordered domains (RGG1 and RGG2 domain) of FUS that leads to an organized and tighter RNA binding. Irrespective of the length of RGG2, the RGG2-RNA interaction is confined to the stem-loop junction and the proximal stem regions. On the other hand, the RGG1-RNA interactions are primarily with the longer RNA stem. We find that the C-terminus of RRM, which make up the “boundary residues” that connect the folded RRM with the long disordered RGG2 stretch of the protein, plays a critical role in RNA binding with the RRM domain. Our study provides high-resolution molecular insights into the FUS-RNA interactions and forms the basis for understanding the molecular origins of full-length FUS interaction with RNA.

## I. INTRODUCTION

FET family of RNA binidng protein aggregate in Amyotrophic lateral sclerosis (ALS) and frontotemporal lobe degeneration (FTLD)^1,2^ two common neurodegenerative diseases usually affecting individuals over 50 years of age. Familial cases of ALS and FTLD contain point mutations^3–5^ in the low complexity regions of FET RBPs^6–8^, which leads to disruption of RNA and protein homeostasis, a major pathogenic mechanism responsible for causing these diseases^9–11^. RNA binding interfaces of RBPs are low complexity regions that are disordered in nature that impart structural flexibility or disorderliness which is an integral part of biomolecular recognition in protein-protein or protein-nucleic acid complexes^12^. Upon RNA binding the low complexity regions transition from disorder-to-ordered state^13^. Fused in Sarcoma (FUS) protein is one such multi-domain protein in the FET family with self-association and RNA binding properties^14–16^. It is present in both nuclear and cytoplasmic biomolecular condensates and plays a key role in RNA metabolism including splicing and transcription. Mutations in FUS cause dysregulation of RNA metabolism and cytoplasmic inclusion, a key event in FUS-associated ALS/FTLD pathogenesis^17^.

FUS binds promiscuously with a wide variety of structured and unstructured RNA and DNA sequences involved in transcription, splicing and DNA repair^18^. FUS is present at high concentrations in the nucleus, yet only 1% of the total concentration is found in nuclear condensates. This phenomenon implies that the phase separation of FUS is dependent on RNA concentration, and a high RNA/protein ratio is reported to prevent phase separation, while a low ratio promotes phase separation^19^. Another study by Hamad et al. using fragments of promoter-associated non-coding RNA reveals RNA sequence-dependent regulation of FUS condensate formation^20,21^. Together, it is clear that phase separation of FUS depends on the concentration of both specific and non-specific RNA. Such an ambiguous behavior can only be explained by the conformational plasticity of the disordered regions of FUS making them adaptive to bind different RNAs. In general, the RNA sequence-dependent interaction and conformational changes in the aggregate-prone disordered regions of FUS protein are hypothesised to be responsible for the regulation of condensate or membrane-less organelles (MLO) formation, although the exact molecular mechanism of this interaction is not well understood.

As shown in Fig. 1(a), FUS is a 526 amino-acids (AA) long protein comprising a low-complexity region enriched with Serine, Tyrosine, Glycine, and Glutamine residues (SYGQ) at its N-terminal (1-165 AA), ordered RNA recognition motif (RRM, 281-377 AA) and zinc-finger (419-454 AA) domains, separated by three Arg-Gly-Gly rich, RGG (RGG2: 378-418 AA, RGG3: 455-501 AA) domains. The region 166-269 AA can be further classified into a G-rich (166-222 AA) and RG/RGG rich (223-268 AA) region, alternatively the entire region from 166-269 is also called RGG1. The NES (269-280 AA) and C-terminal PY-NLS (502-526 AA) regions help in their cytoplasmic and nuclear localization^22^. There is an ambiguity in defining the boundary between RRM and RGG2 domains involving the residues 360-SGNPIKVSFATRRADFNR-377. An important outcome of our study is the important role played by these boundary residues, which we discuss in detail in our paper. The RNA binding regions in FUS are the folded RRM and ZnF domains along with the three disordered RG/RGG-rich regions. The N-terminal SYGQ domain (also called the low complexity domain, LCD) is primarily responsible for the phase separation and aggregation behavior of FUS. The predominant occurrence of SYGQ residues and their arrangement in the protein sequence are the essential determinants of FUS self-assembly propensity. Several studies have elucidated the importance of aromatic repeats and the importance of their arrangement towards the phase separation in IDPs including FUS^23,24^. Computational studies have played a major role in understanding the molecular interactions among LCDs, in particular the contributions of Arginine and Tyrosine residues towards regulating the liquid/gel/solid states of FUS^25^. Apart from the N-terminal LCD, the FUS protein contains three RG/RGG-rich disordered regions with nucleic acid binding ability that is also known to mediate phase separation^25,26^. Recently, inter/intra-molecular interactions between LCD and RGG regions have been identified as another driving force in stabilizing the FUS condensates^25^.

**FIG. 1:**
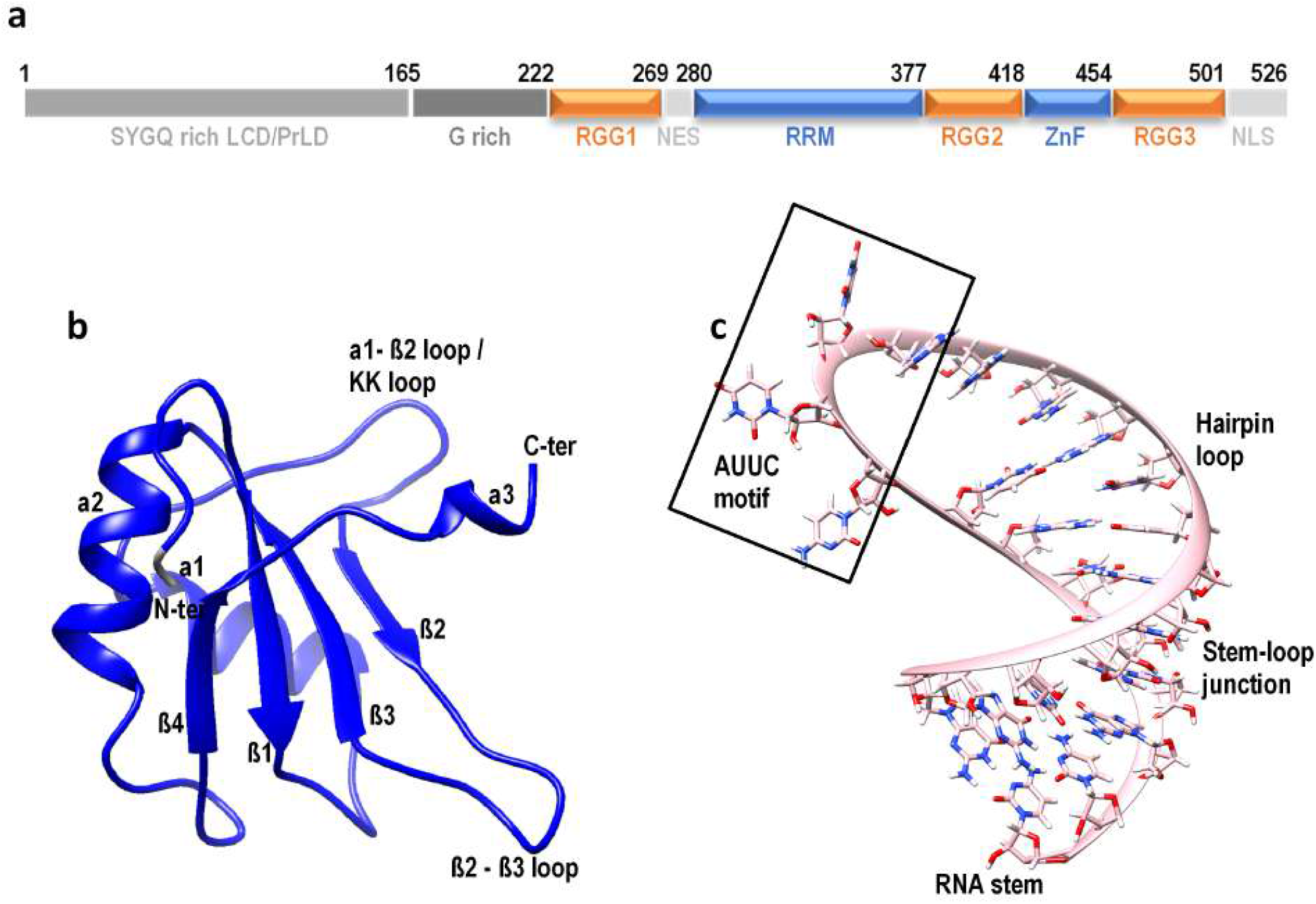
The domain organization of FUS. The RRM and ZnF (in blue) are the only folded domains, while the RGG (in orange) are disordered regions rich in RG/RGG motifs with nucleic acid binding properties. The three-dimensional structure of (b) RRM domain and (c) RNA stem-loop structure with marked secondary structures motifs.

The RRM domain of FUS^27^ is a folded domain, known to recognize several RNA as well as DNA targets in the genome, and multiple pieces of evidence exist for its recognition of a wide range of RNA and DNA structures^28,29^. The RRM domain comprises *β*1 − *α*1 − *β*2 − *β*3 − *α*2 − *β*4 fold with a single short helical turn at the C-terminus (structure shown in Fig. 1(b). The RNA binding pocket includes the surfaces of *β*-sheets 1, 2, and 3, the *α*1-*β*2 hairpin loop (also called KK loop) conserved in the FET family proteins, the *β*2-*β*3 loop and the C-terminal helical turn. A previous docking study of RRM with a 12mer ssRNA has established the RNA-binding importance of the loop dynamics^30^. RNA recognition by FUS-RRM is mainly driven by the positively charged residues due to the lack of aromatic amino acids over the *β*-sheet surface and the longer *β*-hairpin connecting *α*1 and *β*2, which is unique and distinct from a canonical RRM^27^. Several studies have identified sequence and structural motifs in RNA that are recognized by FUS^28,31^. The widely known RNA sequence motifs are GGUG, CGCGC and GUGGU, while the structural motifs are an AU-rich stem-loop structure (Fig. 1c)^28^, and a G-quadruplex structure^32^. A recent NMR study by Loughlin et al.^22^ has identified the structure of the RRM domain in complex with a stem-loop structured hnRNP A2/B1 pre-mRNA (Fig. 1c). This study claims a shape specificity for the RRM domain and identifies a consensus motif of “NYNY” (N=Cyt/Ura/Ade/Gua; Y=Cyt/Ura) sequence in the single-stranded loop of the stem-loop RNA as the recognition motif. The FUS ZnF domain is another ordered nucleic acid binding domain in FUS, which shows specificity for a GGU motif. The NMR structure of the ZnF domain in complex with a 5mer RNA of sequence UGGUG has been solved by Loughlin et al. to establish the binding mode and specificity of the ZnF domain. Together with the sequence specificity of the RRM domain, Loughlin et al. propose the recognition of a bipartite motif in a stem-loop RNA (YNY and GG[U/G] within a 30 nt separation) by the RRM-RGG2-ZnF construct of FUS expressing both shape and sequence specificities.

The binding affinity of different domains of FUS with RNA has been identified previously by Jacob Schwartz and co-workers^26^. This isothermal titration calorimetry study has shown that all FUS domains express weak binding affinity with RNA when present individually^26,29^. The binding affinity of wildtype FUS is 0.7 *µ*M, while the two folded domains, RRM (*>* 90 *µ*M) and ZnF (*>* 175 *µ*M) show very weak affinity individually. Among the three disordered RGG regions, the RGG1 with 3 *µ*M is the strongest, followed by RGG3 with 9 *µ*M and RGG2 with 61 *µ*M. However, when the two weak binding domains RRM and RGG2 are present together, the binding affinity shows a drastic increase to 2.5 *µ*M. This is further enhanced to 1.9 *µ*M when RGG1 is also included. Such a major jump in binding affinity among the individual (RRM and RGG2 with *>* 90*µ*M and 61 *µ*M, respectively) and combined RRM-RGG2 (2.5 *µ*MM) constructs clearly implies cooperativity between these folded and disordered regions to bind RNA, which has not been investigated yet. Our study analyzes the interaction of the RRM domain with RNA and explores the possibility of a cooperative RNA binding mechanism between RRM and RGG2 through all-atom molecular dynamics simulations. Though the importance of FUS-RNA interaction has been well elucidated, the details of molecular interactions at the single molecule level are still lacking. In this context, our study finds merit in exploring the characteristics of FUS-RNA interaction from the perspective of a varying number of RGG repeats. It is previously established that the RGG regions interact with LCD in a condensate. Together with our observations of RGG-RNA interactions, it is possible that there are RNA-mediated interactions between LCD and RGG in a condensate. Hence, our study forms the basis for addressing an interesting mechanistic hypothesis regarding the RNA concentration-dependent phase behavior of FUS condensates.

The rest of the paper is organized as follows. We describe our modeling and analysis methods in detail in the “Material and Method” section. Besides providing information on the molecular simulation protocols and reporting the systems under consideration, we also provide details about how we reconstructed these RNA-protein complexes with IDPs flanking on both sides of the folded RRM region. We have also used some ingenious approaches to analyze our complex trajectory data and we also describe that in this section. In the section after this, which is the Results and Discussion section, we highlight our salient findings. We find that the C-terminal helix in the RRM-RGG2 boundary region weakly holds together the RRM-RNA complex and the flanking RGG domains play a major role in enhancing RNA binding, with a number of repeats of RGG coming across as a major factor in the stable RNP complex formation. We also show how the sequence and length of the RNA are important in these complexes. We close the paper with a short conclusion section.

## II. MATERIALS AND METHOD

### A. Molecular dynamics simulations

Molecular dynamics (MD) simulations of the FUS-RNA complexes were performed in the GROMACS package using a99SB-disp forcefield for Protein^33^ and OL3 forcefield for RNA^34^. The various FUS-RNA complex systems under consideration are listed in Table I. The forcefield a99SB-disp has been used successfully in recent years to sample proteins containing both folded and disordered regions, and hence this was used in our study. These complexes were solvated with TIP4P-D water specific for a99SB-disp in a periodic box with an additional water pad extending up to 12 Å in all directions. The systems were neutralized and additional ions were added to mimic a salt concentration of 150 mM. The short-range interactions were truncated with a cut-off distance of 10 Å. Electrostatic interactions were treated by particle-mesh Ewald with a real space cut-off value of 10 Å. Bonds containing hydrogens were constrained using the LINCS algorithm. The solvated and neutralized systems were energy minimized using the Steepest Descent algorithm followed by equilibration of 5 ns period and subsequently, production runs were taken up after assuring stable equilibration. The temperature and pressure of the systems were maintained at 310 K and 1 Atm using the Nose-Hoover thermostat and Parrinello-Rahman barostat in an NPT ensemble. The simulations were performed in triplicate of 500 ns each to improve sampling and the significance of our results. All analyses, besides the ones described in the subsections below, were performed with the Gromacs analysis tools and CPPTRAJ module of AmberTools20.UCSF Chimera v1.13 and VMD 1.9.3 were used for visualization and preparing the images.

### B. Modeling RNA stem-loop structure

The structure of a stem-loop RNA formed by the hnRNP A2/B1 pre-mRNA sequence solved in complex with the FUS-RRM domain by NMR (PDB ID: 6GBM^22^) was used in our study. Other FUS-RNA complexes were modeled using this 23mer stem-loop RNA by superposing the RRM domains. To extend the length of this RNA, the hnRNP A2/B1 pre-mRNA sequence with the bipartite motif (RRM specific AUUC and ZnF specific GGU) was used. This sequence, used by Loughlin et al.^22^ has an RNA hairpin with a single-stranded stem. The extended RNA structure was modeled as a double strand by extending the complementary strand also in order to use a stable RNA structure while modeling the flexible RGG loops. The RNA structure was modeled using the Discovery studio visualizer 2019 and the *FUS*_418_ complex structures were modeled based on the binding orientation of the 23mer RNA hairpin in 6GBM by superimposing the RRM domains. The 23mer RNA hairpin has the following sequence: GGCAGAUUACAAUUCUAUUUGCC. The following sequence was used for the bipartite motif used by Loughlin and co-workers^22^ [GAUUAGGU-UUUGUGAGUAGACAGAUUACAAUUCUAUUUUAA] and we use an extended sequence as described above and given as: [GAUUAGGUUUUGUGAGUAGACAGAUUACAAUUCUAUUUGUCUACUCACAAAACCUAAUC]

### C. Modeling of RGG1 and RGG2 stretches

Computational modeling of IDP and IDR structures^35,36^ is a challenging process due to their heterogeneous conformations landscape. Also, IDRs that follow the “folding upon binding” principle generally require their interaction partners to attain a properly folded state. There are several integrative modeling and pure simulations methods, both at all-atom resolutions and reduced resolutions, which can be used to elucidate the conformational ensemble of IDP/IDRs in their APO state^37–50^. In our study, where the IDR needs to be modeled in complex with the RNA, we add the IDR in fragments and have modeled the RGG repeats undergoing the “folding upon binding” mechanism using classical all-atom molecular dynamics simulation of the interacting partners. The RGG regions were added sequentially to the RRM (PDB ID: 6SNJ^51^). In other words, the RGG2 was first added to the RRM-RNA construct, and following this, the RGG1 was modeled into the system. The sequence of RGG2 (391-418 AA) and RGG1 (223-269 AA) were split into fragments of 3-5 AA (8 and 7 fragments for RGG2 and RGG1, respectively) and each fragment was added one at a time. After adding each fragment, the rest of the FUS domains, including the RNA (RRM-RNA for RGG2 modeling and RRM-RGG2-RNA for RGG1 modeling) were restrained to the initial position with a harmonic restraint weight of 10000 KJ and simulated for a period of 50 ns. After the 50 ns restrained simulation, the trajectories were analyzed for their interaction with RNA while the C-(for RGG2) or N-(for RGG1) terminal residues are in an extended state to allow further extension. The snapshots matching these criteria were extracted from the trajectory and another fragment was added to this structure to repeat the 50 ns restrained simulation. This procedure was repeated until all fragments were added. Following this, all harmonic restraints were removed and the structures were simulated using the standard Molecular Dynamics simulation protocol as explained above.

### D. Interaction analysis

The inter-atomic distance maps representing the distance between each pair of residues were calculated as an average of the last 100 ns of one of the trajectories. Since the RRM domain is quite stable, our discussions are confined to the inter-molecular distances between FUS and RNA. Hence, the distance maps were plotted with FUS on the x-axis and RNA on the y-axis with a distance cut-off of 20 Å. This distance ensures that all interacting residues and interaction types including electrostatic, *π*-, Hydrogen bonds, and hydrophobic interactions are accounted for during the calculation. The amino acid-wise interaction plots were calculated as an average of the last 100 ns of all three independent simulations. Cpptraj module of AmberTools20 was used to extract all pairs of residues between FUS and RNA present within a 6 Å distance that is maintained for at least 10% of the simulation period. The interactions by each RNA base were clustered on the interacting amino acids and the number of these interactions is plotted. Since all residue pairs within 6 Å are considered, the obtained number includes all types of non-bonded interactions like hydrogen bonds, electrostatic, *π*- and hydrophobic interactions. The interactions were classified based on similar studies done previously^52^.

### E. Uniform clustering of simulation IDR-RNA ensemble using t-SNE

Molecular dynamics simulation generates an ensemble of conformations representing the dynamics of biomolecules and valuable insights could be derived by clustering these conformations. The clustering of an IDP ensemble is a challenging task due to the high conformational heterogeneity. Several clustering methods like hierarchal, vector quantization and neural network are available to perform the clustering analysis. In our study, we use the nonlinear dimensionality reduction method called t-distributed Stochastic Neighbor Embedding (t-SNE) coupled with the k-Means method for clustering the highly heterogeneous IDP/IDR ensemble of FUS into subgroups of homogeneous conformations. Complete details about this method for clustering IDPs are available in the recent paper from our group^53^. The clustering was driven by calculating the RMSD of every conformation with every other conformation, extracted at 50 ps interval, to represent the similarity/dissimilarity among the ensemble. The RMSD was calculated for the RGG2 region while superposing the stable RRM domain in order to account for the dynamics of RGG2 alone. The major advantage of t-SNE algorithm is the tunable parameter called perplexity value, which can balance the information between the local and global features of our dataset. The choice of perplexity value is important for dividing the data into discrete and unambiguous clusters. In this work, different perplexity values and the number of K-means clusters were explored and the combination that gave us the best possible Silhouette score was used to undertake the clustering exercise. Our in-house code and SciKit, an open-source library for Python-based machine learning was used to perform these analyses.

Input files needed to initiate molecular simulations and full trajectory data of all simulations for all systems considered in this work are available on our server for download. The server data can be accessed via our laboratory GitHub link: https://github.com/codesrivastavalab/RNA-FUS-AAMD codesrivastavalab/RNA-FUS-AAMD. The files can also be accessed directly from our SharePoint location [here](https://indianinstituteofscience-my.sharepoint.com/:f:/g/personal/bsangeetha_iisc_ac_in/Er0MtllWY-9EnxpAum3fs6MBDYJ_m3JHQzt7Bl4uIF-YyQ?e=RbG79d).

## III. RESULT AND DISCUSSION

### A. Boundary residues between the folded RRM and disordered RGG2 is critical for tight RNA binding

There exists an ambiguity in defining the boundary between the RRM and RGG2 domains. Several reports consider this boundary to be present at different residues in the region 360-377 AA, with a majority of them considering at 371 AA^22 27 54-57^. Hence, we modeled a core RRM-RNA complex (276-368 AA) and the structure is shown in Fig. 2(a). The minimum distance between any pair of atoms among the core RRM and RNA (Fig. 2(b)) showed that the minimum distance remained within 2 Å for the initial 40 ns and starts fluctuating thereafter. The minimum distance increases continuously from 60 ns and after 85 ns, the distance shows a drastic increase indicating the dissociation of RNA from the core RRM (moviefile1.mpeg in SI). The inter-atomic distance matrix also clearly shows the dissociation of RNA from the core RRM as plotted in Fig. 2(c). Hence, it is clear that the core RRM is insufficient to bind the RNA and the residues beyond 369 play an important role. Accordingly, it has been reported previously by Liu et al.,^27^ that a chemical shift perturbation was observed for the residues 369-376 AA upon nucleic acid binding. The presence of these residues (369-ATRRADFNR-376 AA) significantly increases the volume of the RNA binding pocket, as seen through our CASTp binding pocket analysis (Fig. S1 in SI). The volume of the binding pocket increases from 43.5 Å^3^ (for RRM 276-368 AA) to 1021 Å^3^ when the RRM includes 276-377 AA. Therefore, in spite of the ambiguity between different studies, we consider the RRM domain boundary at 377 AA, the minimal region required to bind RNA. Also, the choice of 377 AA is in accordance with previous studies including the NMR structure solution studies by Loughlin et al.^22,57^, the structure used in our study. It is also significant to note that the boundary residues 369-377 AA form a single helical turn-like structure expressing six Hydrogen bonds and two cation-*π* interactions with the RNA in the NMR solution structure (shown in Fig. S2).

**FIG. 2:**
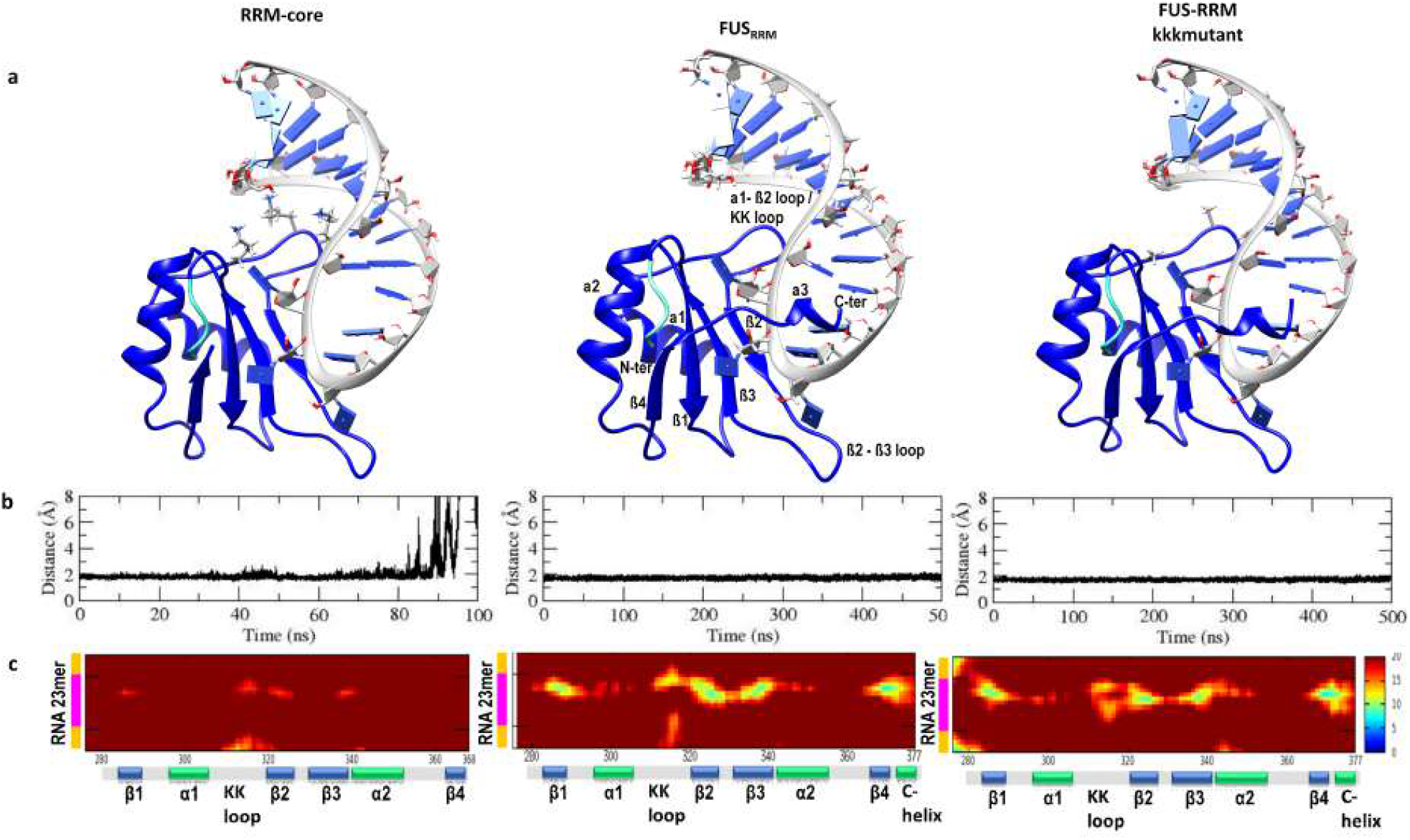
Dynamics of FUS_*RRM*_. (a) The initial structures of RRM-core (276-368 AA), FUS_*RRM*_ (276-377 AA), and KK loop mutant (K312A/K315A/K316A). The NES residues 276-280 are colored in cyan. (b) Variation in the center of the mass distance between the RRM domain and RNA. (c) inter-atomic distances (in *Å*) between the residues of FUS RRM and RNA averaged over the last 100 ns simulation. The secondary structures of RRM are represented on the x-axis, while the RNA stem (yellow) and RNA loop (magenta) are represented on the y-axis.

We simulated the RRM-RNA complex (PDB ID: 6GBM, 276-377 AA, Fig. 2(a)) in triplicates where each replica was run for 500 ns each. We find similar behavior in all three replicates. The root means squared displacement (RMSD) (Fig. S3 (a) in Supporting Information (SI)) shows that the RRM domain is highly stable with an RMSD variation of less than 5 Å. The nature of RNA binding with respect to the stable RRM domain was monitored by calculating the RMSD of RNA as a whole while superposing the RRM. This RMSD indicates the stability of the binding orientation of RNA with respect to RRM, and a large variation up to 15 Å indicates the dynamic and unstable binding of RNA. The distance between the center of mass (com) of the two molecules was monitored in Fig. S3(b), and the variation of about 5 Å indicates a weak/flexible RNA binding. Though the RMSD and com-com distances indicate unstable RNA binding, the minimum distance between any pair of atoms in RRM and RNA lies within 2 Å (Fig. S3(b)) showing that at least parts of RNA remain in contact with the RRM. Hence, to identify the important interacting regions, we monitored the inter-atomic distances between every residue pair in RNA and RRM averaged over the last 100 ns. In Fig. S3 (c), the inter-atomic distances decrease on a Red to Blue scale and brighter intensities depict tighter binding. We observe that the distance between RNA and the C-terminal helix, KK-loop, and *β*3-*α*2 loop stabilizes after simulation, as compared with the interatomic distances of the initial complex (Fig. S3(c)).

Structurally, the interaction of RNA with RRM can be classified based on the interacting regions as (i) the surfaces of *β*-strands 1, 2 and 3 with the recognition motif AUUC, (ii) the *β*2-*β*3 loop with AUUC motif, (iii) the KK loop with the major groove of the stem-loop junction, and (iv) the C-terminal helical turn with RNA backbone (Fig. 1). The superposition of the RRM-RNA complex before and after 500 ns simulation clearly depicts the unwinding of the C-terminal helix and its displacement from the initial position leading to a loss of interactions with the RNA backbone (Fig. S3(d)). Similarly, the displacement of RNA also leads to the disruption of interactions with the KK loop. The interaction of the AUUC recognition motif with different regions of RNA is focused in a 2-dimensional interaction diagram for clarity. In the initial complex, at least 6 hydrogen bonds were formed by the C-terminal helix with RNA (Fig. S2), while several of these were lost after simulation (Fig. S3 (e,f)). Even though the residues Phe288, Arg328, and Lys334 were expressing *π*-stacking or *π*-cation interactions with the RNA bases, the interacting pairs from the initial complex are not conserved after simulation. Altogether, the stem of RNA binds weakly with the RRM domain, yet the RNA motif AUUC expresses several strong contacts with the *β*-sheets of the RRM domain. Moreover, the boundary residues making up the C-terminal helix play a major role in holding the RNA close to the RRM domain.

It is commonly believed that the Lys residues in the KK loop (312, 315 and 316 AA) are important for RNA binding and subcellular localization. Moreover, mutational studies on the KK loop revealed similar chemical shifts for the mutant RRM-RNA/DNA complex and mutant-apo RRM indicating that the mutation impairs nucleic acid binding^27^. However, our simulations have highlighted a significant role for the boundary residues between the RRM and RGG2. In addition to the already know KK loop, this so-far unexplored C-terminal region of RRM (369-377 AA) plays a significant role in stabilizing the RNA. The importance of this C-terminal region for RNA binding has been vastly overlooked to date. Though NMR studies have identified their involvement in RNA binding by NMR chemical shift changes^27^, the KK loop has been mainly attributed to the RNA binding property since it is unique to FUS-RRM. In order to understand the importance of the KK loop for RNA binding, we modeled an RRM-RNA complex with KK loop mutations (K312A/K315A/K316A) as shown in Fig. 2(a). The minimum distance between the RRM and RNA during the 500 ns simulation of the mutant RRM-RNA complex does not show any dissociation of RNA and the distance remains within 2 Åduring the entire 500 ns simulation Fig. 2(b). Though there was no dissociation, the inter-atomic distance matrix clearly shows a distinct pattern of RRM-RNA interaction when compared to the FUS_*RRM*_ complex. (Fig. 2(c)). The distance between the NES (276-280 AA) and RNA stem decreases, while the distance between the KK loop and RNA stem increases. Simultaneously, the distance between the KK loop and RNA hairpin loop decreases indicating a rearrangement of the RNA. Based on these results where we witness several rearrangements in the RNA binding pose leading to weaker binding, we hypothesize that the KK-loop mutation prevents initial recognition and binding of RNA/DNA. This is consistent with the experimental observation that mutation in the unique KK-loop of FUS-RRM impairs or greatly reduces the nucleic acid binding affinity^27^.

Due to the lack of stacking interactions between FUS RRM and RNA, it is reported previously that the stability is driven by electrostatic interactions, mainly contributed by the KK loop. However, our study shows that these electrostatic interactions alone are insufficient to stabilize the RNA in the absence of 369-377 AA. These two regions are positioned to interact with RNA from the opposing sides and together they bind both the grooves of the RNA stem-loop structure. The C-terminal region also extends into the RGG2, which is also reported to possess RNA binding activity. According to biochemical studies carried out by Jacob Schwartz and co-workers, the presence of RGG2 increases the RNA binding affinity of FUS, and the affinity also depends on the number of RGG repeats present^26^. Hence, we further extended our study to include varying lengths of RGG repeats and explore its significance in increasing RNA binding affinity.

### B. Electrostatically dominant RGG2-RNA interaction is modulated by the number of RGG repeats

The RGG2 spans residues 378-418 and has five RGG repeats across this sequence. A previous study by Jacob Schwartz and co-workers established that a minimum of three RGG repeats are required to enhance RRM-RNA affinity, and further addition of RGG repeats enhanced the binding affinity closer to the wild-type range. In order to explore the molecular basis for the enhanced binding affinity when including RGG2, we simulated RRM-RGG2-RNA complexes with a varying number of RGG repeats (listed in Table I) and analyzed their interactions with RNA. Initially, the role of the first three RGG repeats (up to 390 aa) was analyzed since the binding affinity shows a remarkable jump with the inclusion of the third repeat, while the presence of only one (up to 380 AA) and two (up to 385 AA) repeats still behaves similar to RRM alone. The coordinates of RGG2 (PDB ID: 6SNJ shown in Fig. 3(a)) were truncated at 380, 385, or 390 to model the three different complexes with a varying number of RGG repeats and these complexes were simulated for 500 ns in triplicates. Fig. 3(b) depicts the center of mass distance between the RRM domain (276-377 aa) and RNA in RRM-RGG2-RNA complexes containing one (*FUS*_380_), two (*FUS*_385_) and three (*FUS*_390_) RGG repeats. When compared with the com-com distance in the RRM-RNA complex, the distance fluctuation decreases in the order of *FUS*_*RRM*_ *> FUS*_380_ *> FUS*_385_ *> FUS*_390_ indicating that the RNA binding is stabilized as the number of RGG repeats increase. And, similar to the RRM-RNA complex, the minimum distance of < 2.2 Å between any residue pair shows that the RNA remains interacting with the RRM domain irrespective of the variation in the com-com distance. Hence, the FUS-RNA interactions were analyzed in detail to understand the interaction of different regions of FUS and the effect of the number of RGG repeats on binding affinities.

**TABLE I.** List of FUS-RNA complex systems studied in this work

| Name | Systems Description | Amino-acids | Structure | Simulation (ns) |
| --- | --- | --- | --- | --- |
| $FUS_{RRM} - core$ | RRM + 23mer RNA | 276-368 | 6GBM | 100 * 1 |
| $FUS_{RRM}$ | RRM + 23mer RNA | 276-377 | 6GBM | 500 * 3 |
| $FUS_{RRM} - KKK_{mut}$ | RRM + 23mer RNA | 276-377 | 6GBM | 500 * 1 |
| $FUS_{380}$ | RRM-RGG2 + 23mer RNA | 260-380 | 6SNJ | 500 * 3 |
| $FUS_{385}$ | RRM-RGG2 + 23mer RNA | 260-385 | 6SNJ | 500 * 3 |
| $FUS_{390}$ | RRM-RGG2 + 23mer RNA | 260-390 | 6SNJ | 500 * 3 |
| $FUS_{418}$ | RRM-RGG2 + 59mer RNA | 260-418 | Modelled | 500 * 3 |
| $FUS_{223-418}$ | RGG1-RRM-RGG2 + mut 59mer RNA | 223-418 | Modelled | 500 * 3 |
| $FUS_{390} - RNA_{mut}$ | RRM-RGG2 + mut 23mer RNA | 260-390 | Modelled | 500 * 1 |
| $FUS_{418} - RNA_{mut}$ | RRM-RGG2 + mut 59mer RNA | 260-418 | Modelled | 500 * 1 |

**FIG. 3:**
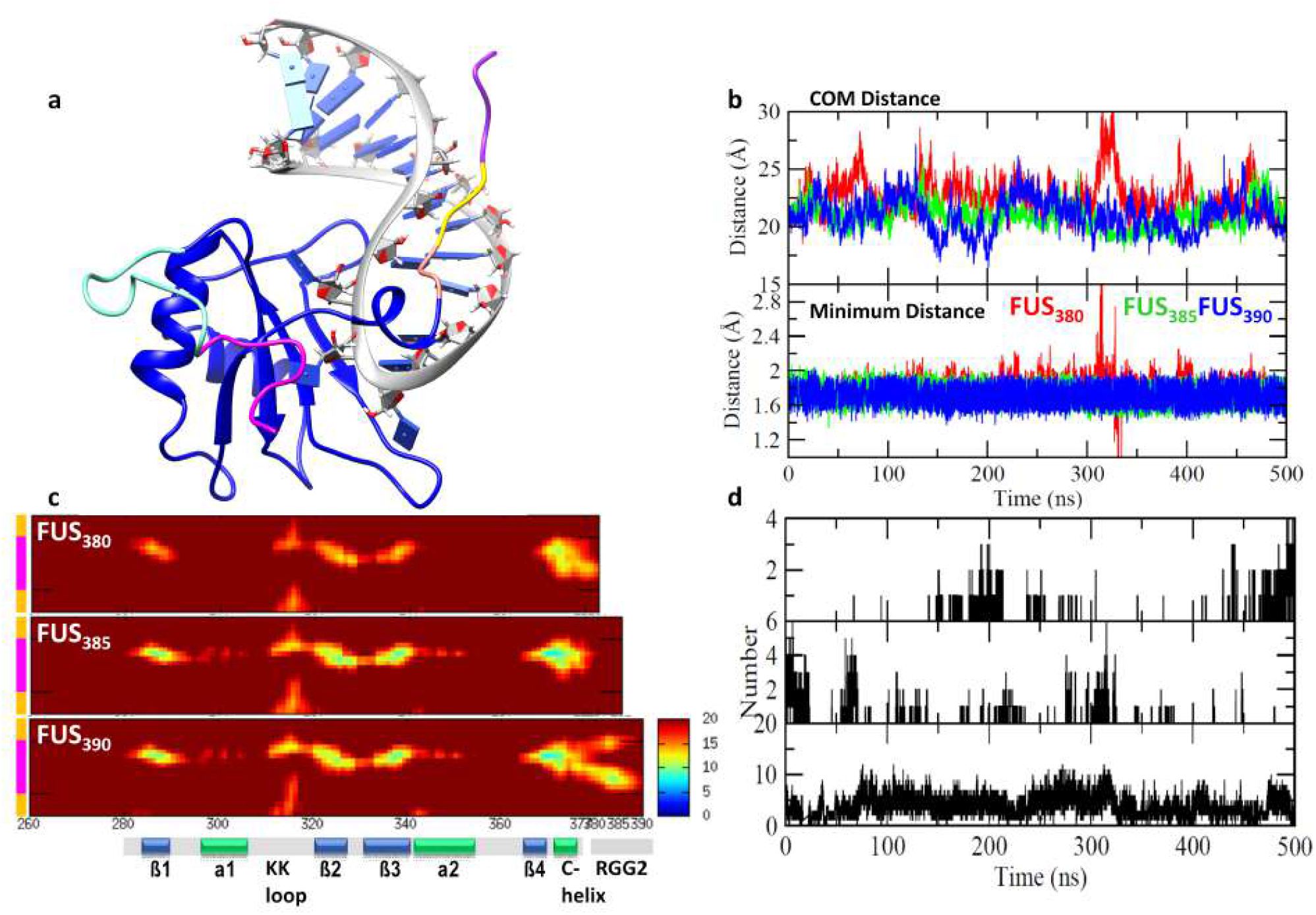
Dynamics of FUS_380_, FUS_385_ and FUS_390_. (a) The structure of FUS_390_ was used to model the truncated structures differentiated as 378-380 AA in Pink, 381-385 AA in Yellow, and 386-390 AA in Purple colors. (b) Variation in the center-of-mass distance between the RRM domain and RNA, and the minimum distance between any pair of atoms in RRM and RNA. (c) inter-atomic distances (in *Å*) between the residues of FUS and RNA averaged over the last 100 ns simulation. The secondary structures of FUS are represented on the x-axis, while the RNA stem (yellow) and RNA loop (magenta) are represented on the y-axis. (d) Time evolution of the number of hydrogen bonds formed between the RGG2 and RNA.

The inter-atomic distance matrices, between the RRM and RNA in the three systems shown in Fig. 3(c) clearly portray the difference arising due to changing lengths of RGG repeats. In *FUS*_380_, the distance between RNA and RRM increases as seen by the reduced intensities for the RNA in general. However, the C-terminal of RRM and RGG2 (370-380 AA) remains close to the RNA. On the other hand, the inter-atomic distance between RRM and RNA in *FUS*_385_ decreases considerably as noted by strong intensities of the RNA loop with the KK loop as well as the *β*-sheets. Notably, the residues 370-380 remain tightly bound to the RNA, while, the residues 381-385 do not express any intensity with the RNA. Upon extending the RGG2 to include the third RGG repeat in *FUS*_390_, the residues 370-390 are closer to the RNA loop and stem-loop junction. The number of H-bonds between RGG2 and RNA also shows an increasing pattern with respect to the increasing number of RGG repeats (Fig. 3(d)). The *FUS*_380_ complex shows ∼ 4 H-bonds at the most, whereas the *FUS*_385_ shows ∼ 2 additional H-bonds. Interestingly, *FUS*_390_ complex expresses the most H-bonds (∼10) between RGG2 and RNA indicating a major drift in the interaction pattern with the addition of only one more RGG repeat.

The C-terminal helix plays a major role in stabilizing the RNA as seen in our previous sections. Visual analysis of the trajectories also reveals interesting changes in the stability of this C-terminal helix and hence we performed secondary structure analysis. Fig. S4 in SI shows that the C-terminal helix is lost in *FUS*_380_, while it is less stable and loses helicity at the end of 500 ns simulation of *FUS*_385_. Interestingly, the C-terminal helix is highly stable in *FUS*_390_, and significantly, the stability of the C-terminal helix has a major influence on the stability of RNA. The complex structure after 500 ns simulation superimposed over the respective initial structures is shown in Fig. S5(a) in SI. The RGG2 in *FUS*_380_ is insufficient to stabilize the RNA, similar to *FUS*_*RRM*_, while in the case of *FUS*_385_, the RGG2 remains coiled near the AUUC motif of RNA. Interestingly, the RGG2 in *FUS*_390_ remains bound to the RNA spine. The structure of RNA in *FUS*_380_ is highly distorted and the RNA loop is pushed out of the binding pocket, which also explains the observed loss of intensities in the inter-atomic distance matrix. Interestingly, the overall RNA structure is conserved in both *FUS*_385_ and *FUS*_390_.

The interaction of the AUUC motif with the RRM domain was monitored in the three systems and the interactions are shown in Fig. S5(b) and Table S1 in SI. Apart from a *π*-interaction with Arg328 and hydrophobic interactions with Tyr325 and Arg372, the RNA in *FUS*_380_ does not show any other interactions with the RRM reinforcing the weak intensities in the inter-atomic distance matrix. Contrarily, the RNA in *FUS*_385_ shows several novel interactions with RRM including *π*-interactions with Thr286, Arg372, and Phe375, H-bond interactions with Tyr325, Thr338, and Arg371, and other hydrophobic interactions. It is noteworthy that in addition to the interactions seen in the NMR complex, the *FUS*_390_ complex shows additional interactions also indicating a tighter binding of RNA.

In order to understand the contribution of various residues in *FUS*_380_, *FUS*_385_ and *FUS*_390_ that interact with each RNA base, we monitored the number of interactions expressed by each amino acid (summing up the electrostatic, hydrophobic and hydrogen bonds). And we present the data as a histogram plot (Fig. S6 in SI). Also, to collectively understand the FUS-RNA interactions in the three independent simulations of each system, the histograms in Fig. S6 were calculated as an average over the last 100 ns of all three trajectories. It is clear from in Fig. S6 that the FUS-RNA interactions are mainly mediated by the RRM domain, while the RGG loop adds only a few interactions to help RNA binding. Also, the interactions between RRM and the RNA loop are dominated by Arg as well as Lys residues. In addition to these, Asp and Phe residues show several interactions over the length of RNA, while the other residues like Thr, Ala, Glu, and Gly express very few interactions. The RNA binding pocket in RRM is lined by three Lysines in the KK loop, one Arginine in the *β*2-*β*3 loop, and two Arginines in the C-terminal helix (see Fig. S7 in SI). In *FUS*_380_, the RNA loop expresses several interactions with Arg, Asp, and Lys residues. However, the RNA binding pocket lacks Asp, apart from one in the C-terminal loop, which points towards a new distinct RNA binding mode. Also, the one Arg residue in RGG2 interacts with the entire RNA loop indicating a very dynamic RNA. It is worth noting that the interaction pattern in *FUS*_385_ and *FUS*_390_ is very similar apart from the interactions involving Arg residues. The Arg contacts in *FUS*_385_ are restricted to the AUUC motif and its flanking bases, contributed entirely by the Arg in the RRM domain. Whereas in *FUS*_390_, the Arg from both RRM and RGG2 are involved in binding the RNA loop and stem-loop junction. Interestingly, all three Arg residues of RGG2 in *FUS*_390_ interact with the stem-loop junction of RNA. This observation clearly highlights that the addition of RGG repeat provides additional interaction sites for RNA, and the role of stabilizing the RNA is shared by both RRM and RGG2. The first RGG repeat is located close to the C-terminal helix in a structurally restrained position to provide any stability to the RNA. Moreover, visualizing the simulation trajectory of *FUS*_380_ also revealed that the C-terminal residues lose their helicity and weaken their interaction with RNA. Further extension of RGG repeats stabilizes the C-terminal helix and mediates their interaction with the RNA as seen in *FUS*_390_.

The RRM domain has five Arginines, three of which line the RNA binding pocket. On the other hand, the RGG2 (377-418 AA) has a compositional bias and contains another five Arginines along with 28 Glycines. The impact experienced by *FUS*_390_-RNA complex over *FUS*_380_ or *FUS*_385_ complexes due to the addition of one RGG repeat was clearly established in the previous section. Hence, we further aimed to explore the reported increase in RNA binding affinity due to the addition of two more Arginines and ∼ 20 more Glycines to *FUS*_390_. The *FUS*_418_-RNA complex was modeled with all five repeats of RGG2 as a highly disordered structure. The modeling was performed by the sequential addition of 3-5 residues with 50ns restrained simulations at each step. In order to accommodate the extended structure, the length of the double-stranded stem of RNA was also extended by adding 10 bp. The modeling protocol is explained in detail in the methods section and the structure of the modeled, as well as the 500ns simulated RRM-RGG2 construct, is shown in Fig. 4(a,b). The residue-specific interaction histogram for *FUS*_418_-RNA complex was calculated in a similar manner to the other RGG2 systems and is shown in Fig. 4(c,d). The interaction pattern in *FUS*_418_ is very similar to the pattern in *FUS*_390_, where the residues Arg and Lys are dominating. Moreover, the number of interactions experienced by Arg of RGG2 with RNA loop and Lys of RRM with RNA stem is higher than in the other systems. In addition to these two residues, Gly of RGG2 also shows several interactions specifically with the stem-loop junction of RNA. Altogether, the interactions in *FUS*_418_ are uniformly spread over the entire length of RNA, namely Arg of RRM and RGG2 with RNA loop, Gly of RGG2 with stem-loop junction, and Lys of RRM with RNA stem, which in turn also maintains the structural integrity of RNA.

**FIG. 4:**
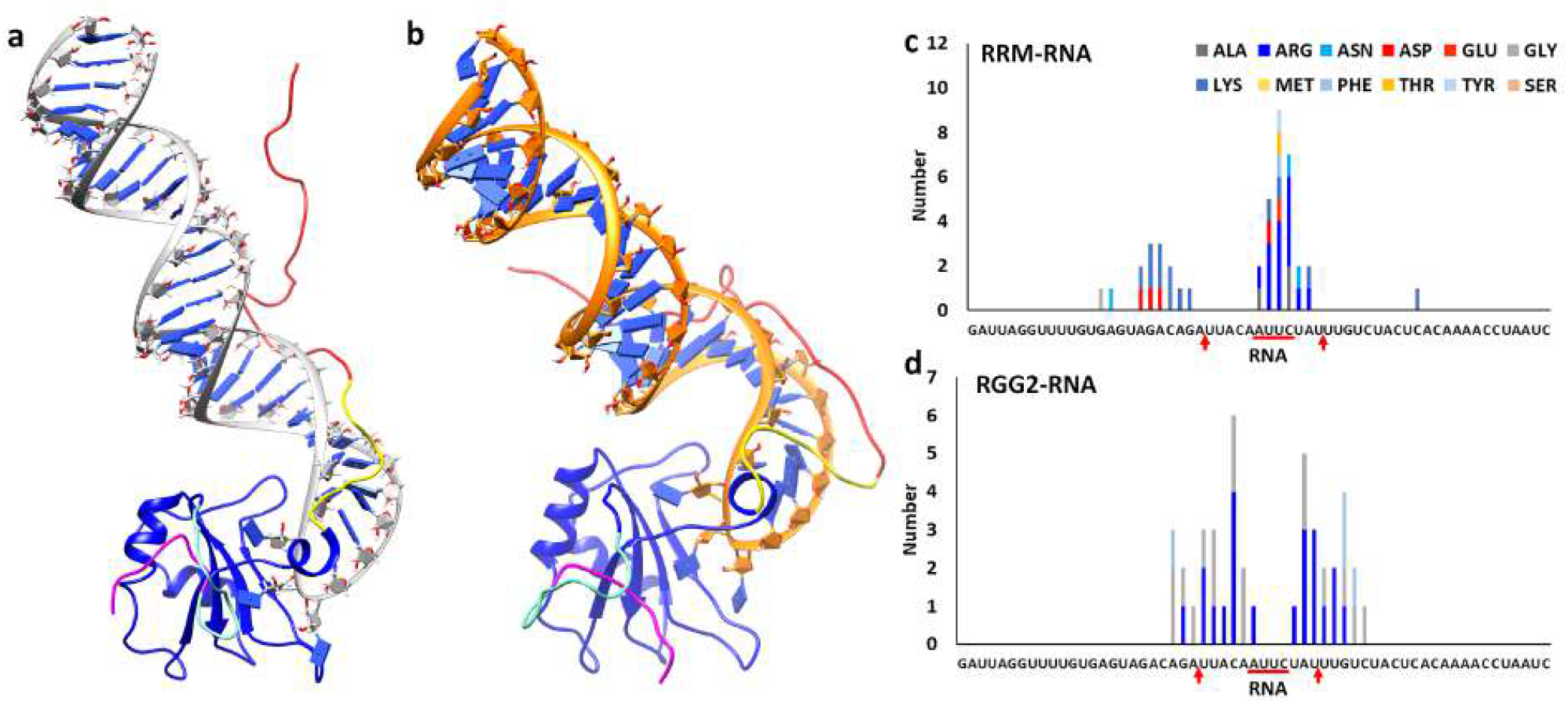
Dynamics of FUS_418_. (a) The modeled structure of FUS-RRM with RGG2 in complex with the 59mer RNA and (b) the 500ns simulated conformation. The different regions of FUS are colored as RGG1 in magenta, NES in cyan, RRM in blue, and RNA in gray (initial) or orange (500ns simulated). The RGG2 up to 390 (yellow) is colored distinctly from 390-418 (salmon) to highlight the importance of this region. Amino acid-wise interactions depicting the number of interactions by each amino acid in the (c) RRM and (d) RGG2 domains with the individual bases of the 59mer RNA.

The contribution of glycine fills the gap in binding the stem-loop junction of RNA (as seen in *FUS*_390_) and vastly enhances the interactions in *FUS*_418_. The nature of these interactions might explain the augmented binding affinity of *FUS*_418_-RNA and hence, together, the interactions are further classified into electrostatic, hydrophobic, and hydrogen bonds (shown in Fig. S8 in SI). The residues Arg, Lys, Asp, and Phe are the major contributors to electrostatic interactions, while the Gly residues are mainly involved in hydrophobic interactions along with a few hydrogen bonds. The major difference between *FUS*_390_ and *FUS*_418_ is the hydrophobic interactions by Glycine stabilizing both strands of the stem-loop junction. Collectively, our study has shown that the increase in the number of RGG repeats has a direct influence on FUS-RNA interactions. It is clear that a large number of strong electrostatic interactions in *FUS*_390_ when compared to *FUS*_385_ might show a greater influence on the binding affinity as reported. However, the comparatively lesser increase in binding affinity between *FUS*_390_ (4.1 *µ*M) and *FUS*_418_ (2.5 *µ*M) is due to the addition of weak hydrophobic interactions by the ∼ 28 Gly residues in RGG2. Though they are weak compared to the electrostatic interactions by Arg and Lys, collectively they might be responsible for the increase in RNA affinity of *FUS*_418_ over *FUS*_390_.

Our *FUS*_418_ simulation allows us to understand the conformational landscape of the structurally less explored RGG2 when interacting with an RNA. The heterogeneous conformations generated in our triplicate simulations were clustered by t-SNE and kMeans methods^53^ to identify the distinct and unique conformations attained by the RGG2. The three-dimensional structures of 10 conformations extracted from each cluster are shown in Fig. 5. It is clearly seen that each cluster is highly homogeneous while the conformations between different clusters are heterogeneous. The conformations of RGG2 were analyzed separately for 378-390 and 391-418 since these two regions show distinct RNA binding behavior. Among the 13 residues in 378-390, at least 52.31 ± 16.25 % of residues remain in contact with the RNA throughout the simulation (% residues in contact with RNA in individual clusters are shown in Fig. 5). On the other hand, only about 31.8 ± 15.9 % of residues among the 28 residues of 391-418 AA are in contact with the RNA. Among the individual clusters, the 378-390 AA shows a consistent interaction with RNA, whereas, the number of residues of 391-418 AA that is in contact with RNA varies widely between 7% to 60%. Altogether, these results clearly highlight that the RGG2 is important for RNA binding and it shows two distinct patterns for RNA binding, stronger binding with < 390 and weaker binding with *>* 390.

**FIG. 5:**
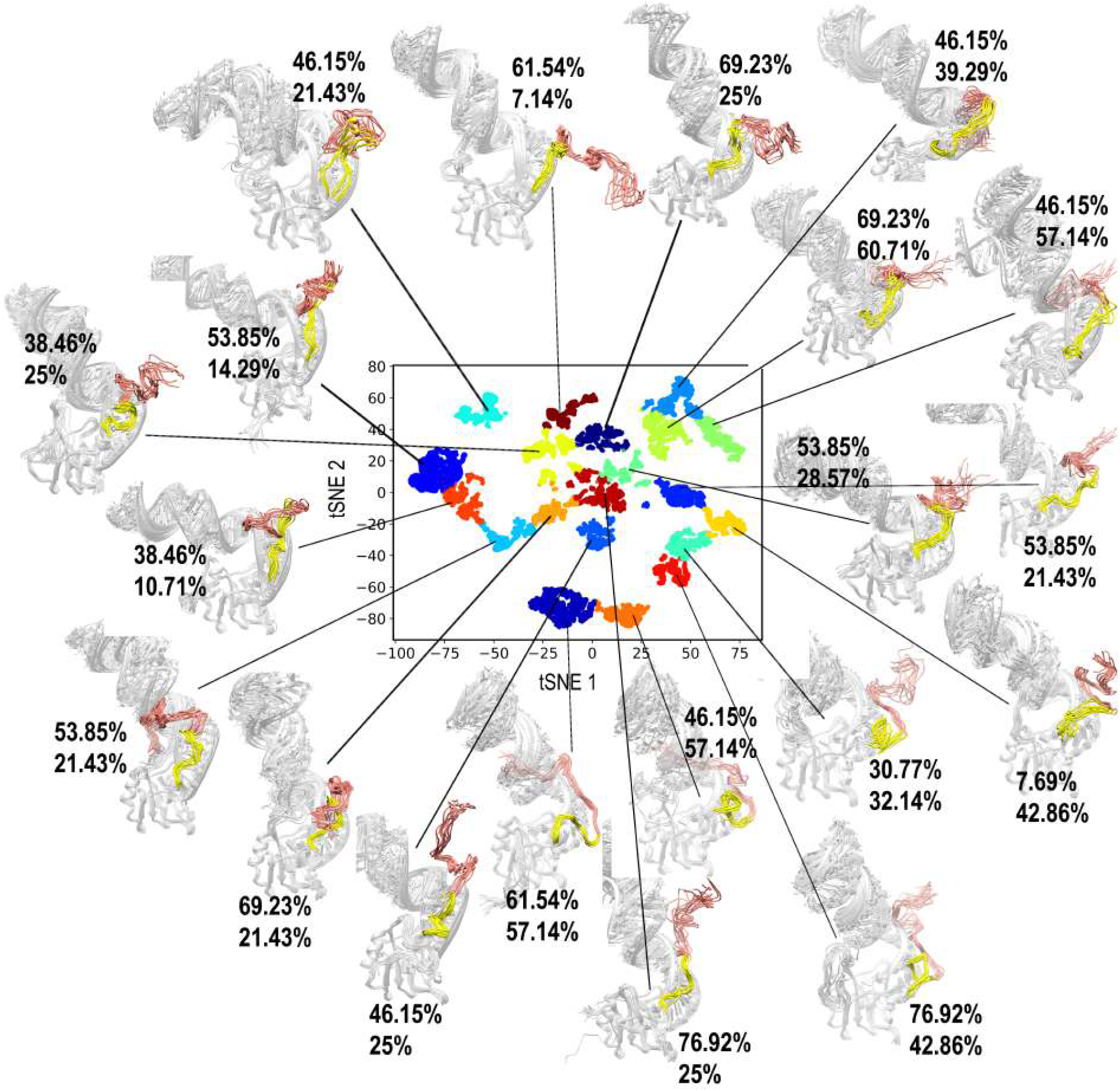
Clustering of the *FUS*_418_ ensemble by t-SNE and kMeans methods. The projection of the first two tSNE components classifies the sampled conformations into 20 distinct and unique clusters. 10 conformers from each cluster are superimposed and the structures are mapped onto the projection. The stable RRM domain and RNA are shown in gray. The RGG2 can be further split into two independent regions (378-390 colored in yellow and 391-418 colored in salmon) based on their interaction pattern with RNA. The percentage of residues in these two regions that are in contact (<3.5 Å) with RNA is marked as % in 378-390 followed by % in 391-418.

### C. Flanking RGGs bind the entire RNA stem and further enhance RNA binding by FUS

The simulation of *FUS*_418_ clearly showed the distinct interaction pattern of RRM and RGG2 with the RNA loop and stem-loop junction, respectively. Even though the longer RGG2 could interact farther on the RNA stem, our analysis has shown that the interactions are confined to the bases close to the stem-loop junction. In particular, the Arg residues in the RGG2 of both *FUS*_390_ and *FUS*_418_ show a very similar interaction pattern with the RNA loop and stem-loop junction, while the additional interactions by the Gly in RGG2 of *FUS*_418_ could be responsible for increasing the binding affinity. Interestingly, these Gly contacts are also limited to the stem-loop junction only, while the farther stem regions remain free of any interactions. Since these interactions saturate at the stem-loop junction, the other regions of FUS should participate to further enhance the RNA binding affinity. Accordingly, the addition of RGG1 (165-267 AA) to the RRM-RGG2 construct is reported to improve the RNA binding affinity to ranges close to wild type. Hence, in order to understand the role of RGG1 in RNA binding, we modeled the RG/RGG rich part of RGG1 (223-267 AA), also in an extended conformation, similar to RGG2. Modeling an additional IDR stretch of ∼ 50 AA to the RRM-RGG2-RNA complex is a non-trivial exercise. The RGG1 was added to one of the clustered conformations of *FUS*_418_ chosen based on the number of residues of RGG1 and RGG2 in contact with the RNA. There are 5 RG/RGG repeats in the 223-267 aa range which might add several interaction sites for the RNA to bind efficiently, and the modeled structure is shown in Fig. 6(a). The RGG1-RRM-RGG2 construct with 59mer RNA, referred hereafter as *FUS*_223*−*418_ was simulated for 500 ns (Fig. 6(b)) and the interatomic distances, as well as the residue-wise interactions, were calculated to understand the FUS-RNA interactions.

**FIG. 6:**
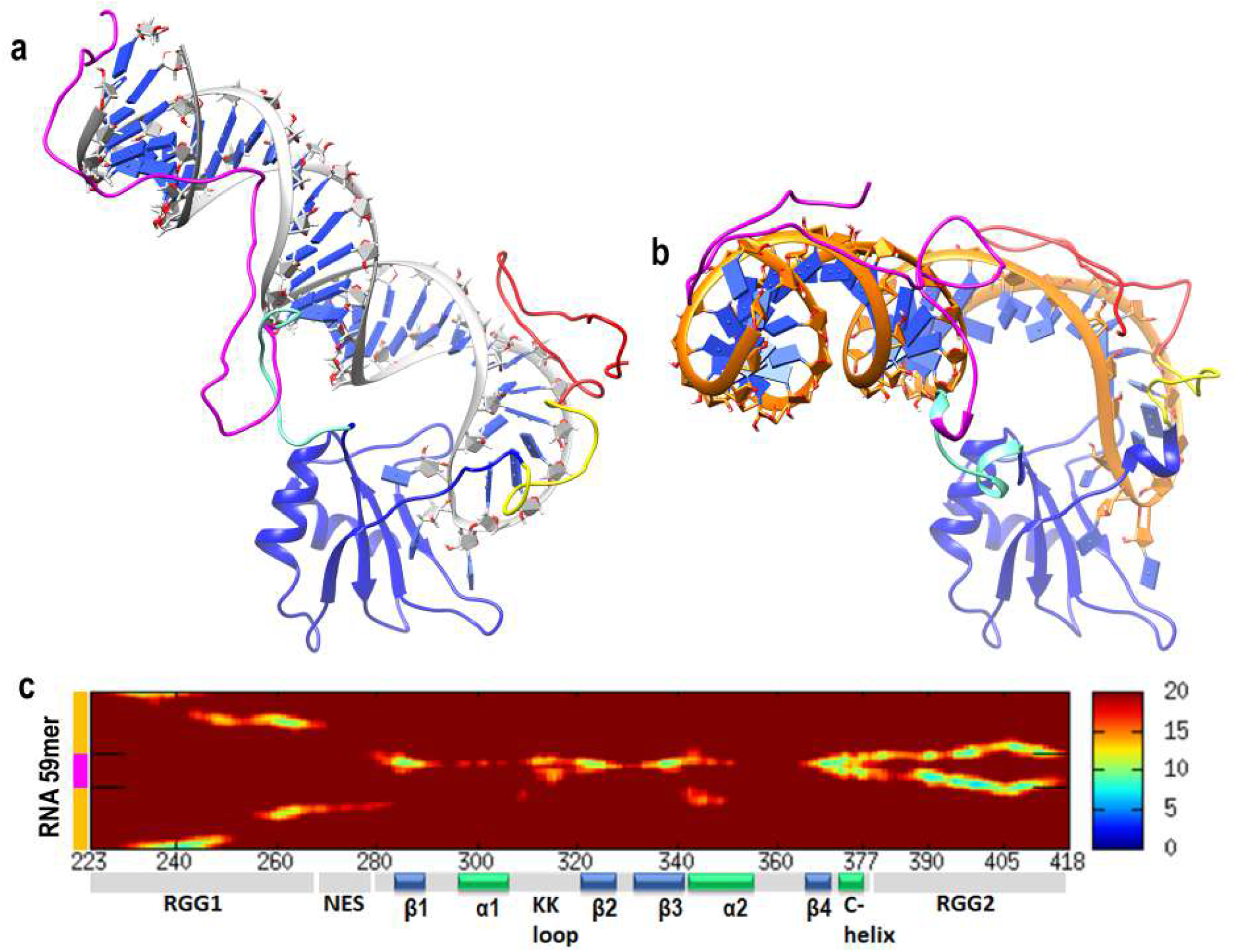
Dynamics of *FUS*_223*−*418_. (a) The modeled structure of FUS-RRM-RGG2 with RGG1 and 59mer RNA and (b) the 500ns simulated conformation. The different regions of FUS are colored as RGG1 in magenta, NES in cyan, RRM in blue, and RNA in gray (initial) or orange (500ns simulated). The RGG2 up to 390 (yellow) is colored distinctly from 390-418 (salmon) to highlight the importance of this region. (c) The inter-molecular distances (in Å) between the residues of FUS (223-418 aa) and RNA averaged over the last 100 ns simulation. The “L” on the y-axis indicates the position of RNA stem-loop junctions.

The inter-atomic distance map in Fig. 6(c) clearly highlights the contacts formed by various regions of RGG1 and RGG2 with the entire length of RNA. The residues of RGG1 remain close to the RNA stem. In particular, the 230-250 AA has high intensity with the ends of the RNA stem. The RRM binds the RNA loop while the RGG2 is strongly in contact with the RNA stem-loop junction. Similarly, the amino acid-wise interactions (shown in Fig. 7), highlight the division of labor by the various domains of FUS to stabilize the RNA by expressing strong electrostatic interactions between their Arg and the RNA. The Arg and Lys residues of RRM interact with the RNA loop, while the Arg residues of RGG2 interact with the stem-loop junction. However, interactions by Gly residues are lesser than *FUS*_418_, which is compensated by the stronger electrostatic interactions by Arg of RGG1 with both the strands of the RNA stem. In addition, Phe, Lys, and Asp also express a few interactions with the RNA stem. Notably, the interactions of RRM and RGG2 with the

**FIG. 7:**
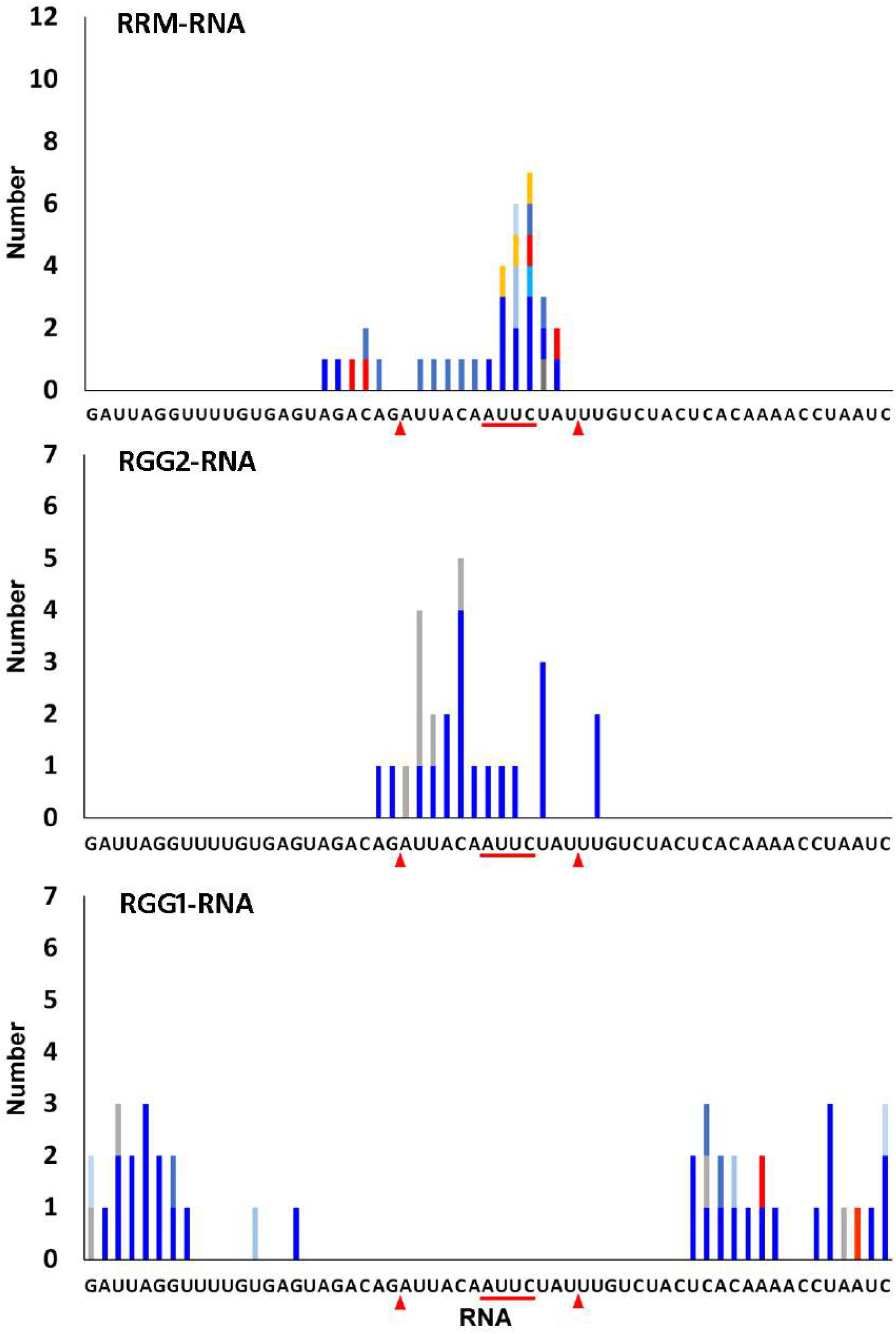
Histogram depicting the number of interactions by each amino acid in the (a) RGG1, (b) RRM, and (d) RGG2 domains with the individual bases of the 59mer RNA of FUS_223*−*418_.

RNA is very similar to those seen in *FUS*_390_ and *FUS*_418_. The three-dimensional structure of the simulated complex is also shown in Fig. 6(b) depicting the wrapping of RGG1 with the double-stranded RNA stem and RGG2 with the spine of the RNA-hairpin. All the sampled conformations of *FUS*_223*−*418_ were clustered by tSNE and kMeans clustering to identify the distinct conformations similar to *FUS*_418_. Due to the highly dynamic behavior of both RNA and RGG1, the conformational landscape is vastly heterogeneous and hence *sim* 70 distinct conformations are populated. In all the 70 clusters, shown in Fig. S9 in SI, the whole of RGG1 is seen to interact with the farther stem of RNA. Altogether, the addition of RGG repeats increases strong electrostatic interactions with RNA, and both the number (*FUS*_390_ vs *FUS*_418_/*FUS*_223*−*418_), as well as the position (RGG1 vs RGG2) of these RGG repeats, have a major influence on the binding of FUS with RNA.

### D. FUS-RRM requires RNA sequence/shape specificity to initiate RNA binding

The RRM domain of FUS is reported to express shape specificity and accordingly, Loughlin et al. proposed a consensus sequence motif of NYNY or YNY (Y=C/U; N=A/G/C/U) for the recognition^22^. The hnRNP A2/B1 pre-mRNA sequence used in our study comprises of AUUC motif at the recognition site and as we saw in the previous sections, this motif interacts well with the RRM domain. For the recognition to happen, the “Y” position in the NYNY motif should contain an “O2” atom as in Cytosine or Uracil. By mutating this position to Adenine or Guanine, we posited that the specificity should be lost and therefore the RRM-RNA interaction should be weaker. In order to test the presence of any sequence or shape specificity in RNA recognition by RRM, we mutated the AUUC motif into AAUG in the NMR structure of *FUS*_390_ and one of the cluster representative structures of *FUS*_418_ (since no structures are reported).

The *FUS*_390_ − *RNA*_*mut*_ and *FUS*_418_ − *RNA*_*mut*_ complexes were modeled and simulated for a period of 500ns and the superimposition of initial and 500ns simulated conformations are shown in Fig. 8 (a,b). The three-dimensional structure clearly shows that the binding of mutant RNA in *FUS*_390_ − *RNA*_*mut*_ is highly disrupted in contrast to the wild-type RNA in *FUS*_390_. Also, the single turn of the C-terminal helix in wildtype RNA complex is extended to include another turn leading to the reorientation of the RGG2 away from the RNA. These major conformational changes were not observed in any of the triplicate trajectories of wild-type RNA complex in *FUS*_390_ suggesting a weak interaction of mutant RNA with RGG2.However, in the case of *FUS*_418_ − *RNA*_*mut*_, the C-terminal helical turn entirely loses its helicity. The inter-atomic distance matrix calculated over the last 100ns of mutant RNA complex simulations is shown in Fig. 8(c,d). The *FUS*_390_ − *RNA*_*mut*_ shows slightly reduced intensities for RRM-RNA and much weaker intensities for RGG2-RNA distances indicating weaker RNA binding. Contrastingly, in the case of *FUS*_418_ − *RNA*_*mut*_, the inter-atomic distance for RRM-RNA is very similar to wildtype RNA complex. Though the 378-390 aa remains close to RNA, the extended RGG2 (391-418 AA) loses interaction with RNA. This is also clear from Fig. 8(d) where the intensities are entirely absent for the extended RGG2 region.

**FIG. 8:**
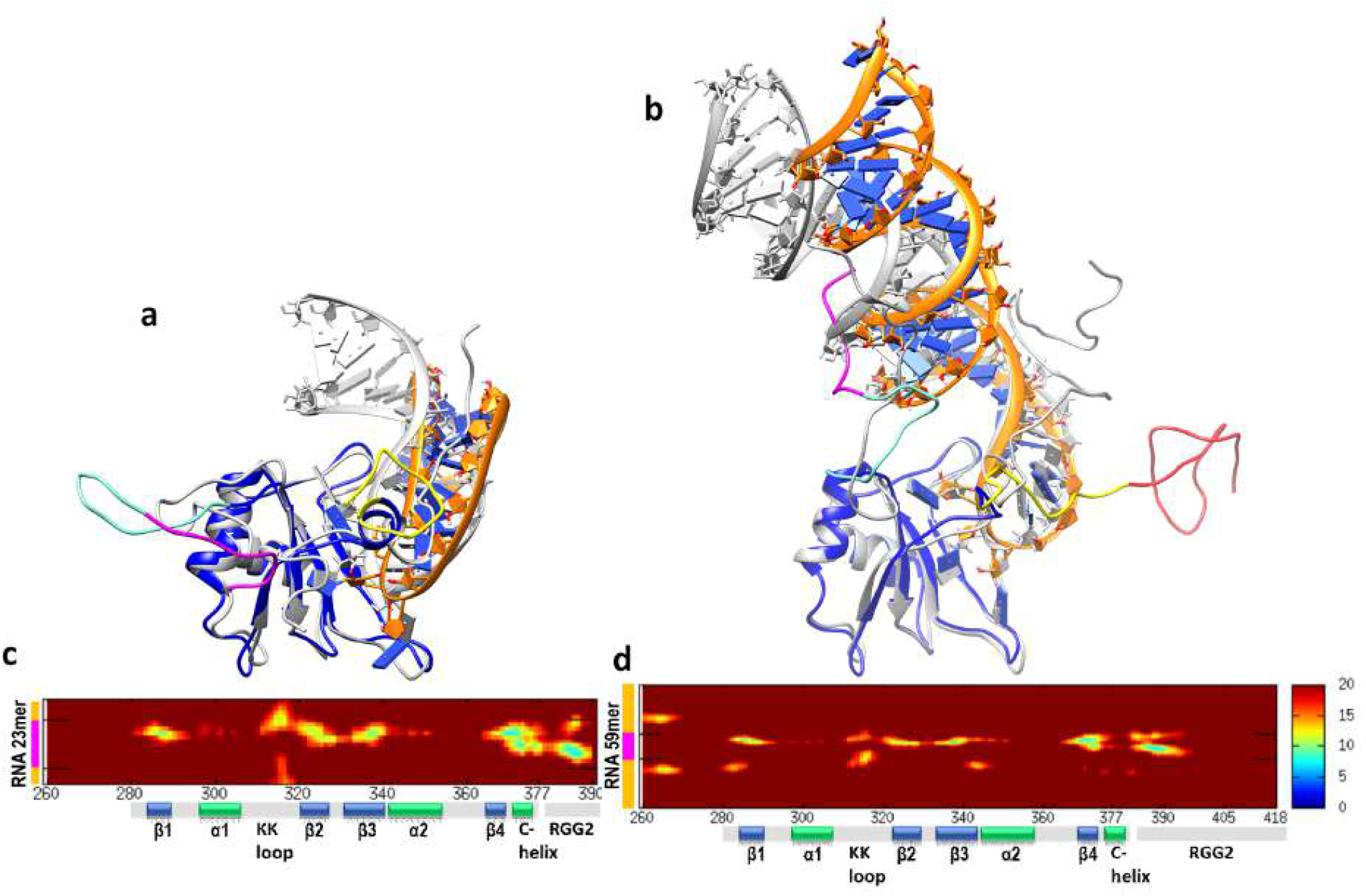
Dynamics of FUS in complex with RNA mutant (a) Structure superposition of initial (gray) and 500 ns simulated conformations of *FUS* − _390_ *RNA*_*mut*_ and *FUS* − _418_ *RNA*_*mut*_. The different regions of FUS in the simulated conformations are colored as RGG1 in magenta, NES in cyan, RRM in blue and RNA in orange. The RGG2 up to 390 (yellow) is colored distinctly from 390-418 (salmon) to highlight the importance of this region. The inter-molecular distances (in Å) between FUS and RNA in (c) *FUS* − _390_ *RNA*_*mut*_, (d) *FUS* − _418_ *RNA*_*mut*_ averaged over the last 100 ns simulation. The “L” on the y-axis indicates the position of the RNA stem-loop

The two-dimensional interaction diagram of the mutated AAUG motif in *FUS*_390_ − *RNA*_*mut*_ and *FUS*_418_ − *RNA*_*mut*_ shows few conserved and several new interactions with the RRM domain (Fig. 9(a,b) and Table S1). The mutation of U in the second position to A allows several additional interactions to form in both the mutant complexes, while none of the interactions from *FUS*_*RRM*_ or NMR are conserved for C to G in the fourth position. The stacking interaction of U in the 3rd position with Phe288 is still conserved along with backbone hydrogen bonds with Thr370 and Arg372. Apart from this, there are several new interactions with the *β*2-*β*3 loop (residues Asn323, Tyr325, and Arg328), C-terminal helix (Thr370, Arg372, and Ala373), and Arg386 of RGG2. When compared with the wildtype complexes, it is clear that the RNA orientation in both the mutant complexes is different and the KK loop is entirely devoid of any interactions. Our results from previous sections have highlighted the importance of hydrophobic interactions by the Gly residues of the extended RGG2 to stabilize RNA. Hence these interactions were further analyzed in *FUS*_418_ − *RNA*_*mut*_, to explore the importance of the extended RGG2 on RNA binding. The interaction histogram of mutant RNA with RRM and RGG2 of *FUS*_418_ − *RNA*_*mut*_ system detailing the contribution of each amino acid type to RNA binding is shown in Fig. 9(c,d).

**FIG. 9:**
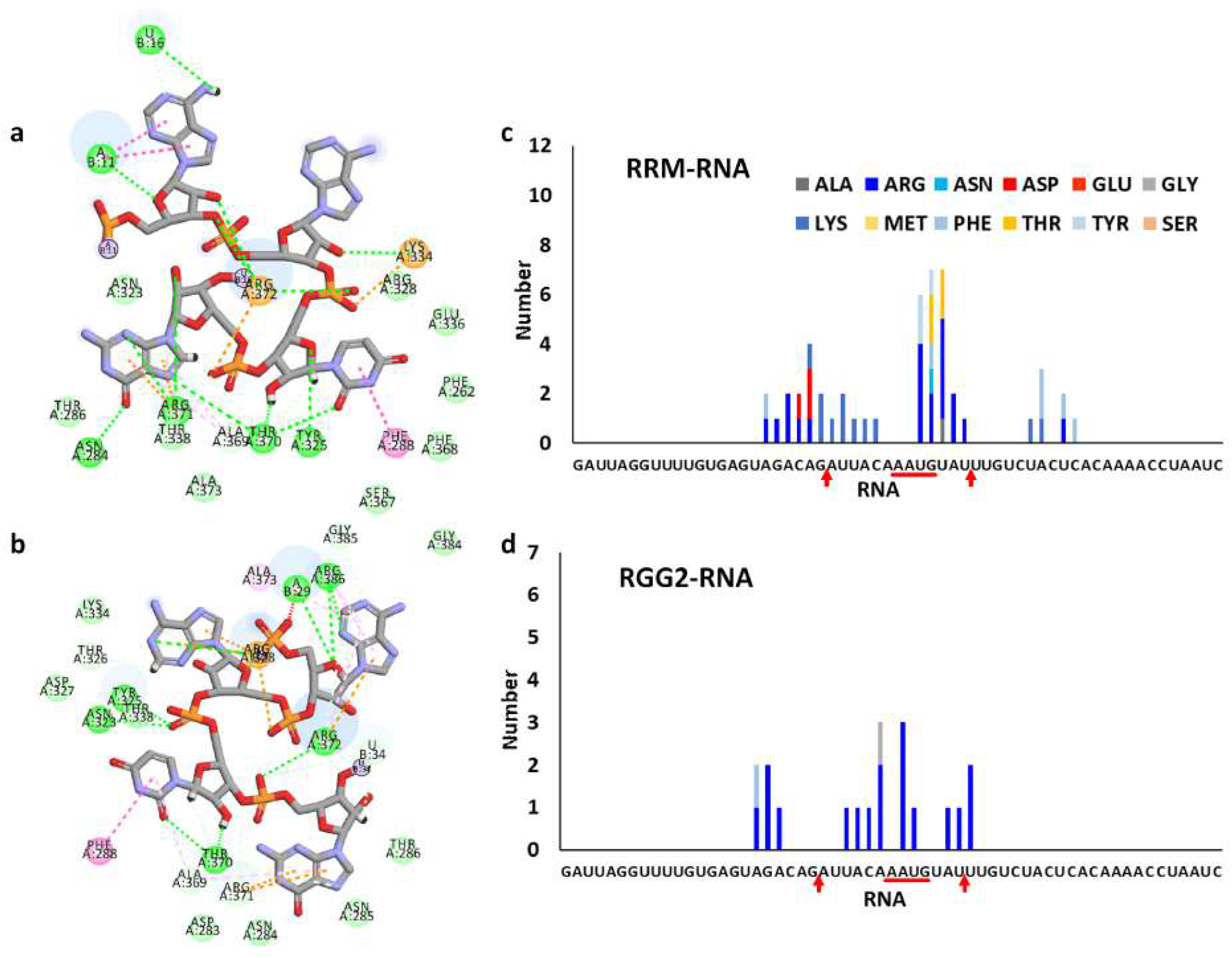
The two-dimensional interaction diagram depicting the different residues interacting with the AUUC motif of RNA in (a) *FUS* − _390_ *RNA*_*mut*_ and (b) *FUS* − _418_ *RNA*_*mut*_. Green dotted lines: Hydrogen bonds, orange dotted lines: *π*-cation interactions, pink dotted lines: *π*-stacking interactions, pale green discs: hydrophobic interactions. Amino acid-wise interactions depicting the number of interactions by each amino acid in the (c) RRM and (d) RGG domains with the individual bases of the 59mer RNA in *FUS*_418_ − *RNA*_*mut*_.

It is surprising that the mutation in the RNA motif recognized by the RRM domain clearly affects the binding and pattern of the remote interactions in the RGG2 loop, particularly with the extended RGG2 (391-418 AA) and its Gly residues. Firstly, the Gly residues are not involved except for only one interaction. Several new interactions between Arg of RRM domain and the RNA stem-loop junction as well as the RNA stem are seen, which is very unique to the mutant RNA complex. Similarly, another unique interaction is seen between the RNA loop and Lys residues. It is worth mentioning here that the *β*-sheet surfaces, where the RNA loop is supposed to interact, are entirely devoid of Lys residues apart from the loops (KK loop and *β*2-*β*3 loop). Even though *FUS*_*RRM*_ was unable to stably bind the RNA, the recognition motif was interacting strongly. However, in the case of the AAUG RNA mutant, the recognition motif loses several interactions with the *β*-sheets of RRM. The loss of these interactions with the RRM is clearly seen to be compensated by stronger electrostatic interactions with the Arg/Lys of both RRM and RGG2. Hence, it is clear that the interaction of RRM with the recognition motif in the mutant RNA loop is severely disrupted, nevertheless, parts of RGG2 were able to hold the RNA stem to still remain interacting with the FUS.

These observations also highlight the importance of RGG2 for RNA binding and the drastic enhancement of binding affinity due to the inclusion of only three RGG repeats. Collectively, it can be hypothesized that the specificity of RRM to RNA sequence/shape is required only for the initial recognition or localization, and thereafter, the interactions with RGG are stronger to overcome any loss in sequence/shape specificity. This indicates a division of labor among the various regions of FUS protein, where the loss of interaction with one of the domains might be compensated by the gain of interactions with the other domains of FUS.

## IV. CONCLUSION

In this paper, we have used large-scale molecular simulations at an all-atom resolution to understand the atomic-level interactions in FUS-RNA complexes. Our study provides molecular-level mechanistic insights into observations from biochemical studies and it has also illuminated our understanding of molecular driving forces that mediate the structure, stability, and interaction of RRM and RGG domains of FUS with a stem-loop junction RNA. Our simulation data clearly brings forth the very important role of the c-terminal region at the interface of RRM and RGG2, which seems to be central to the fidelity of the complex. This region is ambiguously classified as either RRM or RGG2 causing inconsistency in comparing the binding affinities among various experimental literature. We show that excluding this region in RRM leads to dissociation of the RRM-RNA complex and this is an experimentally testable hypothesis. With FUS-RRM devoid of the classical recognition motifs seen in the FET family, we believe that this boundary region between the folded and disordered domains gains importance as the anchor along with the earlier discovered non-canonical central KK loop. Our study also provides the structure biophysical rationale for why at least three RGG repeats are required in RGG to improve binding to the RNA. We find that the Arg residues in the first two RGG repeats are sterically hindered from structurally accessing the RNA due to the persistence caused by the small helix at the start of RGG2. We also find that whatever the length of RGG2 is, the interactions are confined to the RNA loop and stem-loop junction only. The fourth and fifth RGG repeats in RGG2 do not significantly improve the binding strength. However, the increase that was observed could be attributed to the hydrophobic interactions between Glycines and RNA. The role of Glycine residues in biomolecular interactions has been widely overlooked, yet we have observed an important role in these interactions. This is interesting from the point of view of bounds put on RGG repeats that maximizes their functional role. On the other hand, we find that once RGG1 is introduced from the N-terminal end of the RRM, RNA binding noticeably increases again.

Flanking RGGs bind the entire RNA stem and our simulations provide a very clear picture of the origins of the enhanced interactions. Interestingly, the NES region connecting RGG1 with RRM does not express any interactions with the RNA. Our data from RNA mutation simulations again provide experimentally testable hypotheses to establish the RNA sequence and structure specificity of FUS protein where we see specificity for NYNY motif. Mutation directly alters the RRM-NYNY interaction pattern and as a result, we observe an indirect allosteric effect in the RGG2. Such adaptable interactions of FUS is mainly responsible for its promiscuous nucleic acid binding property and minimal sequence specificity.

## V. AUTHOR CONTRIBUTIONS

S.M. and A.S. conceived the research. A.S. and S.B. designed the various simulation experiments in consultation with S.M.; S.B. generated all the trajectories and performed all the calculations; S.B. analyzed the generated data with help from S.M. and A.S.; S.B. wrote the paper with contributions from S.M. and A.S.

## VI. ACKNOWLEDGMENTS

A.S. thanks the Department of Science and Technology (DST) for the National Supercomputing Mission grant (DST/NSM/R&D HPC Applications/2021/03.10) and the Supercomputing Education and Research Center (SERC) at IISc-Bangalore for the high-performance computing (HPC) resources. S.B would like to acknowledge financial support from the NSM grant. The high-performance computing facility “Beagle” setup from grants by a partnership between the Department of Biotechnology of India and the Indian Institute of Science (IISc-DBT partnership program) is greatly acknowledged. A.S. and S.B. also acknowledge the uninterrupted access to the computing facility from the Sankhyasutra HPC enterprises and the consumable grant from the IISc-DBT partnership that supported the access to the facility. S.M would like to thank IISc-Bangalore and the Ministry of Education, India for her start-up grant and the DBT-Wellcome Trust India Alliance for the Intermediate Fellowship. All authors thank Ashok Sekhar for critically reading and commenting on the manuscript.

## Supporting Information

**FIG. S1:**
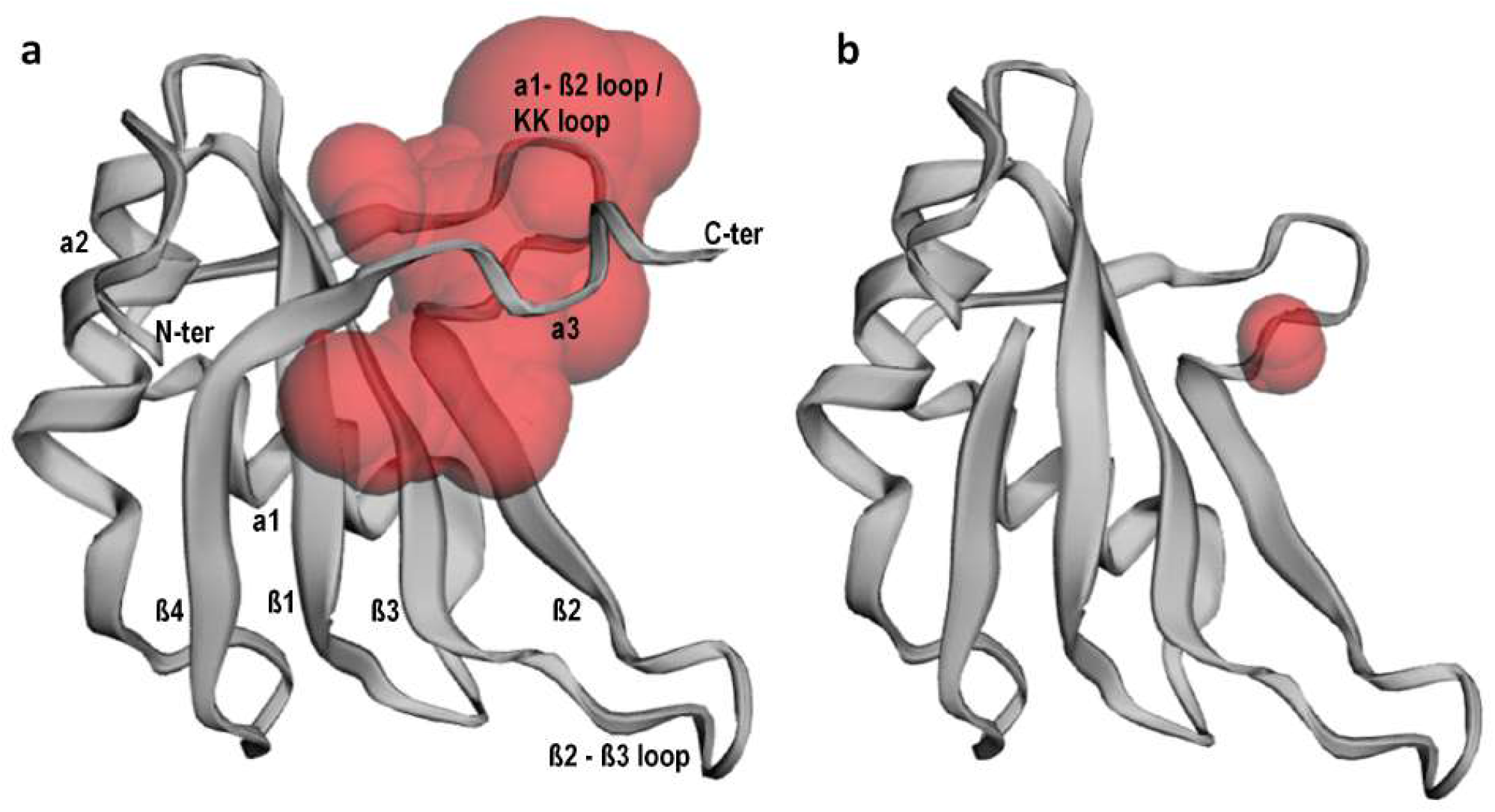
Binding pocket volume analysis using the CASTp server for (a) FUS-RRM (276-377 aa) and (b) truncated FUS-RRM (276-368 aa) to depict the role of the C-terminal helical turn in increasing the volume of RNA binding pocket (red surface representation).

**FIG. S2:**
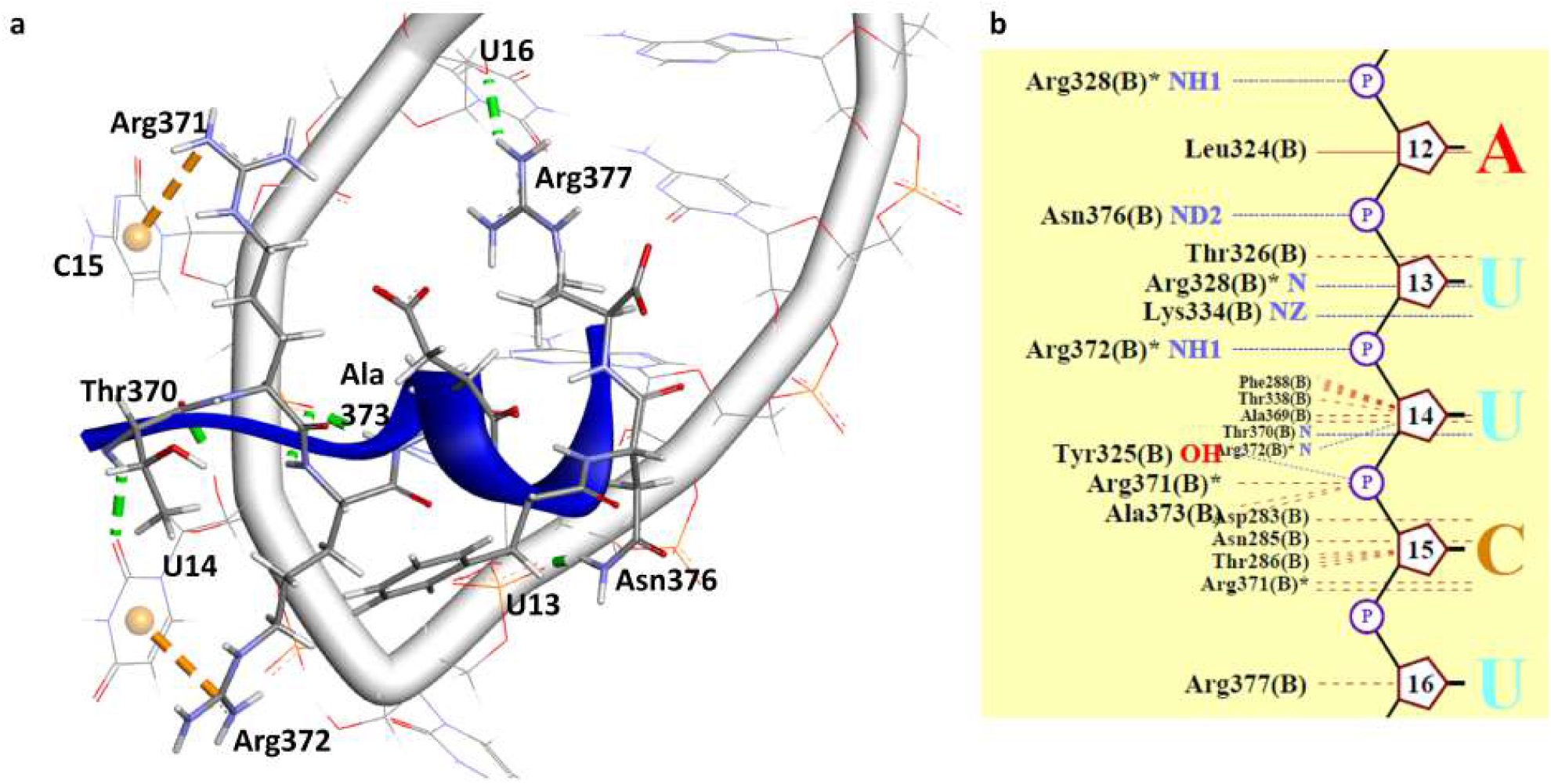
Interaction of C-terminal boundary region (369-377 AA) with RRM in the NMR structure with PDB ID: 6GBM in (a) three-dimensional and (b) two-dimensional representations. FUS is shown in blue cartoon with the sidechains depicted in licorice, colored based on the atoms. The RNA backbone is shown as a tube with the bases displayed as wires. The Hydrogen bonds are shown as green dotted lines, while the *π*-interactions are shown as orange dotted lines. The FUS residues and RNA bases involved in these interactions are labeled. In the two-dimensional representation, blue dotted lines indicate Hydrogen bond and red dashed lines indicate all nonbonded contacts within 3.35 Å.

**FIG. S3:**
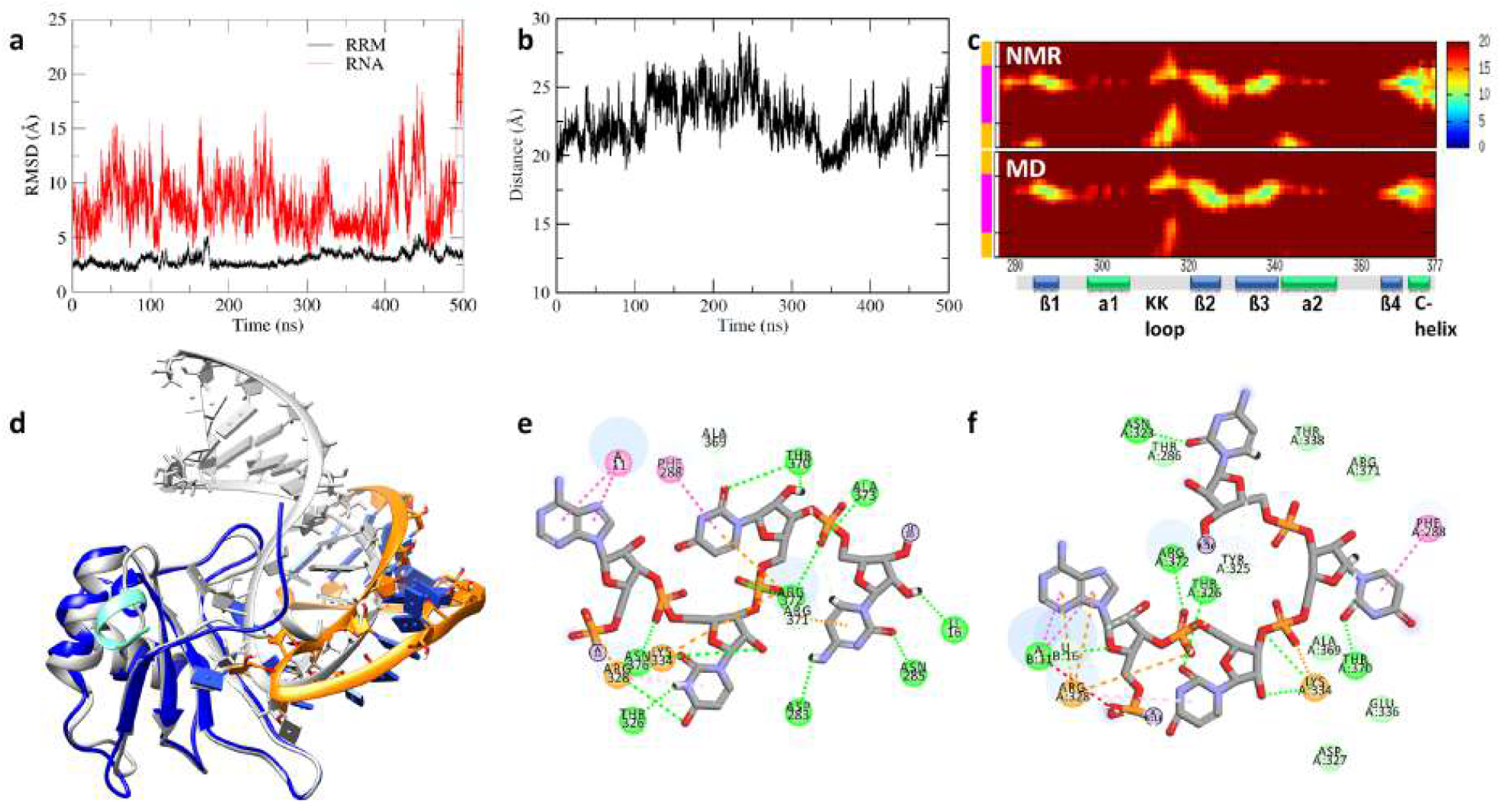
Dynamics of *FUS*_*RRM*_. (a) Time evolution of all-atom RMSD of *FUS*_*RRM*_ (black) and RMSD of RNA (red) calculated with respect to the RRM domain as reference defining the stability of the RNA binding pose. (b) Variation in the center of mass distance as well as the minimum atom-pair distance between the RRM domain and RNA. (c) The inter-atomic distances between the residues of *FUS*_*RRM*_ (276-377 aa) and RNA averaged over the last 100 ns simulation. The “L” on the y-axis indicates the position of RNA stem-loop junctions. (d) Structure superposition of initial (gray) and 500 ns simulated conformations of *FUS*_*RRM*_. The different regions of FUS in the simulated conformations are colored as NES in cyan, RRM in blue, and RNA is colored in orange. The two-dimensional interaction diagram depicts the different residues interacting with the AUUC motif of RNA. Green dotted lines: Hydrogen bonds, orange dotted lines: *π*-cation interactions, pink dotted lines: *π*-stacking interactions, pale green discs: hydrophobic interactions.

**FIG. S4:**
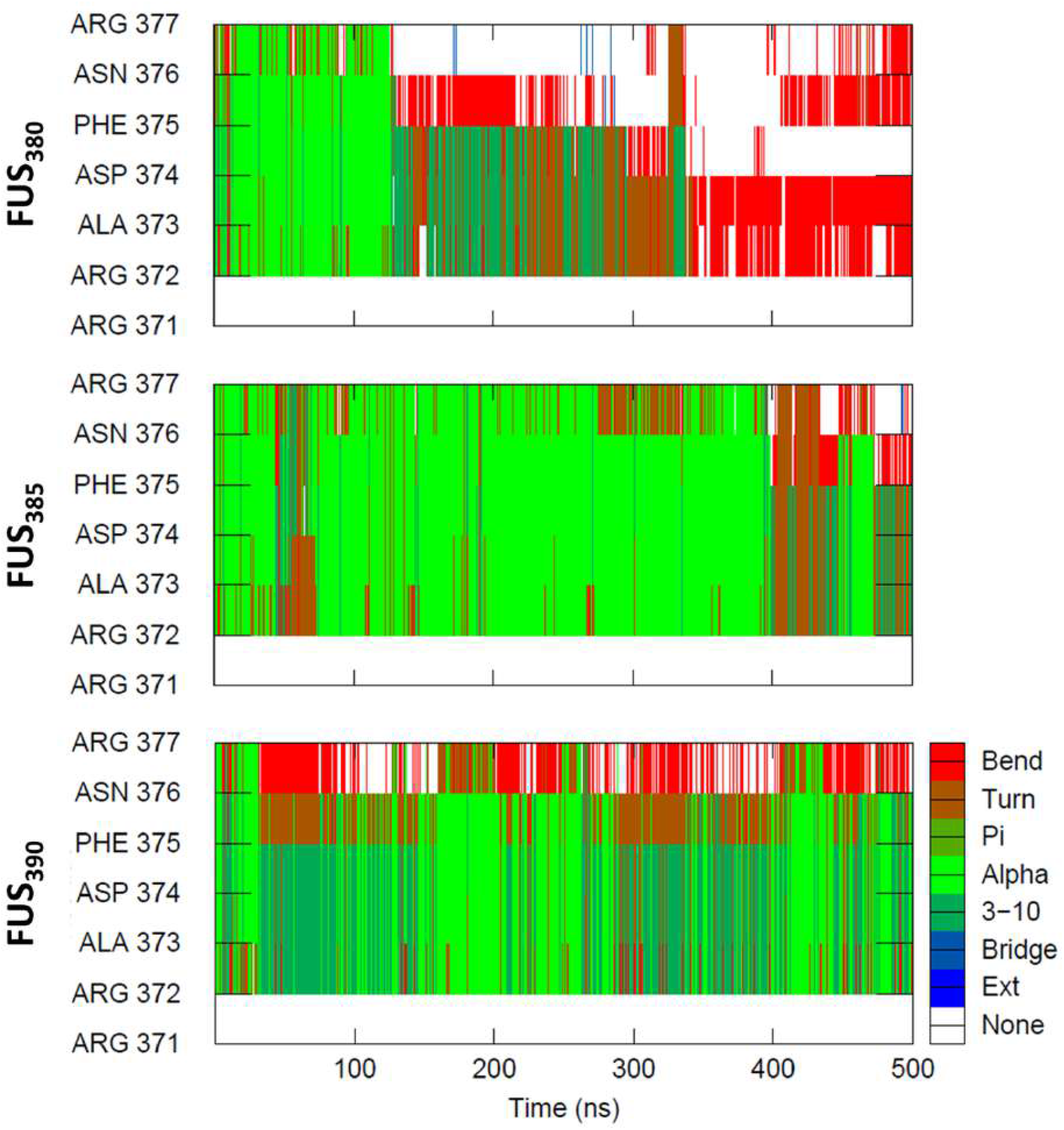
Secondary structure analysis depicting the stability of the C-terminal helix in *FUS*_380_, *FUS*_385_, and *FUS*_390_ systems. Light green: *α*-helix, Dark green: 3_10_ helix, Red: turns.

**FIG. S5:**
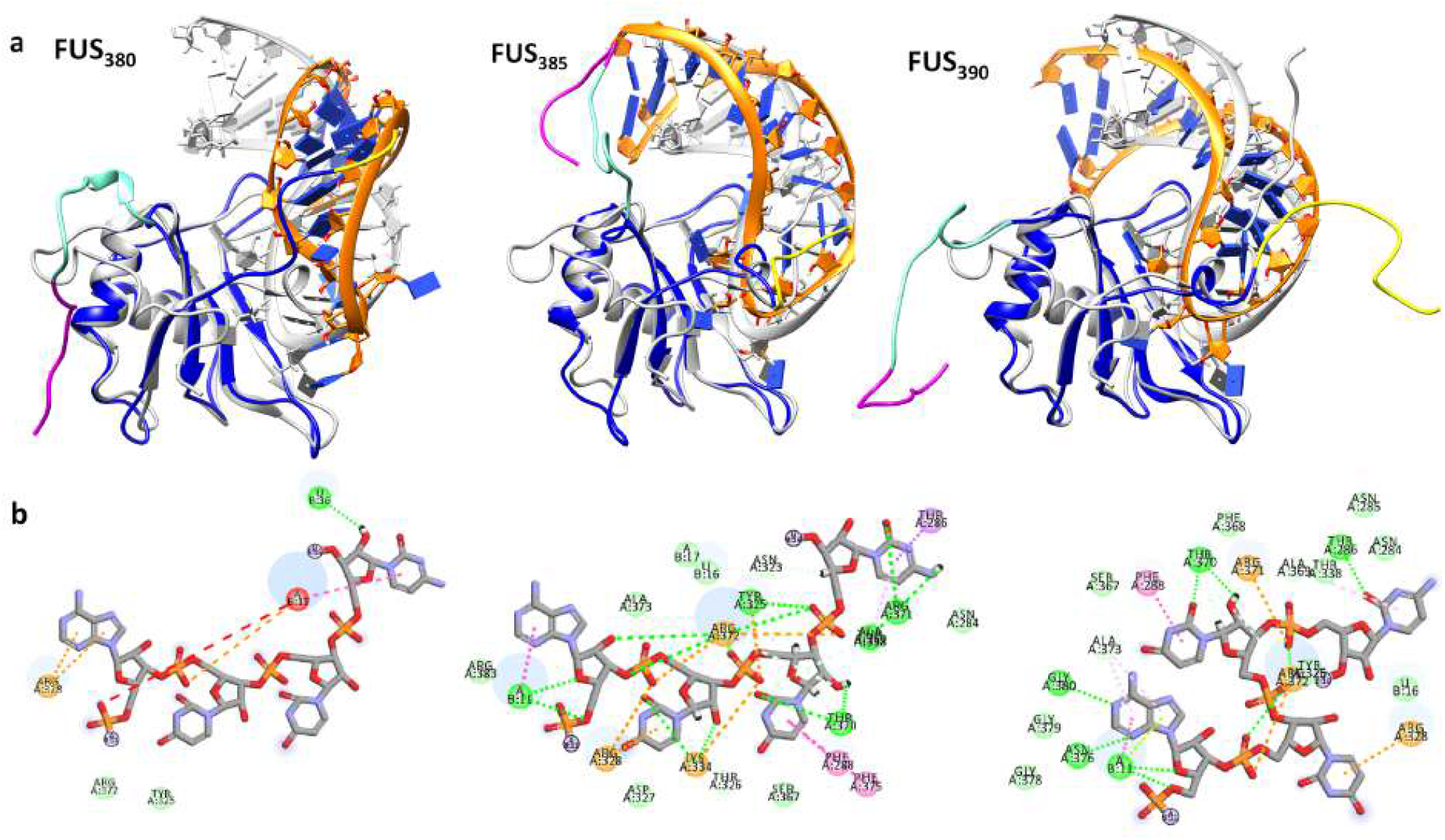
(a) Structure superposition of initial (gray) and 500 ns simulated conformations of *FUS*_380_, *FUS*_385_ and *FUS*_390_. The different regions of FUS in the simulated conformations are colored as RGG1 in magenta, NES in cyan, RRM in blue, and RGG2 in yellow, while the RNA is colored in orange. (b) The two-dimensional interaction diagram depicting the different residues interacting with the AUUC motif of RNA in *FUS*_380_, *FUS*_385_, and *FUS*_390_. Green dotted lines: Hydrogen bonds, orange dotted lines: *π*-cation interactions, pink dotted lines: *π*-stacking interactions, pale green discs: hydrophobic interactions.

**FIG. S6:**
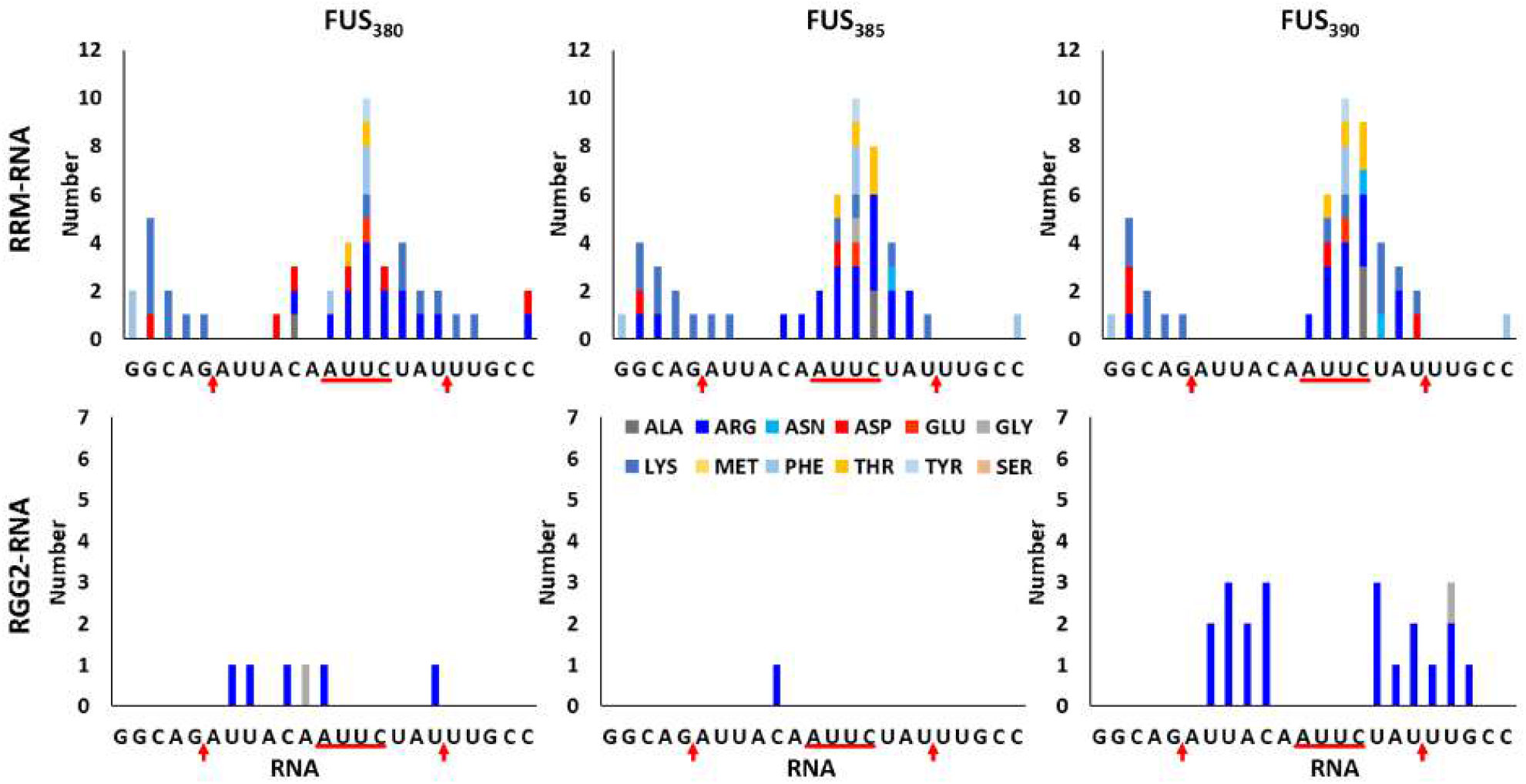
Amino acid-wise interaction histogram depicting the number of interactions by each amino acid in the RRM and RGG2 domains with the individual bases of the 23mer RNA of *FUS*_380_, *FUS*_385_ and *FUS*_390_.

**FIG. S7:**
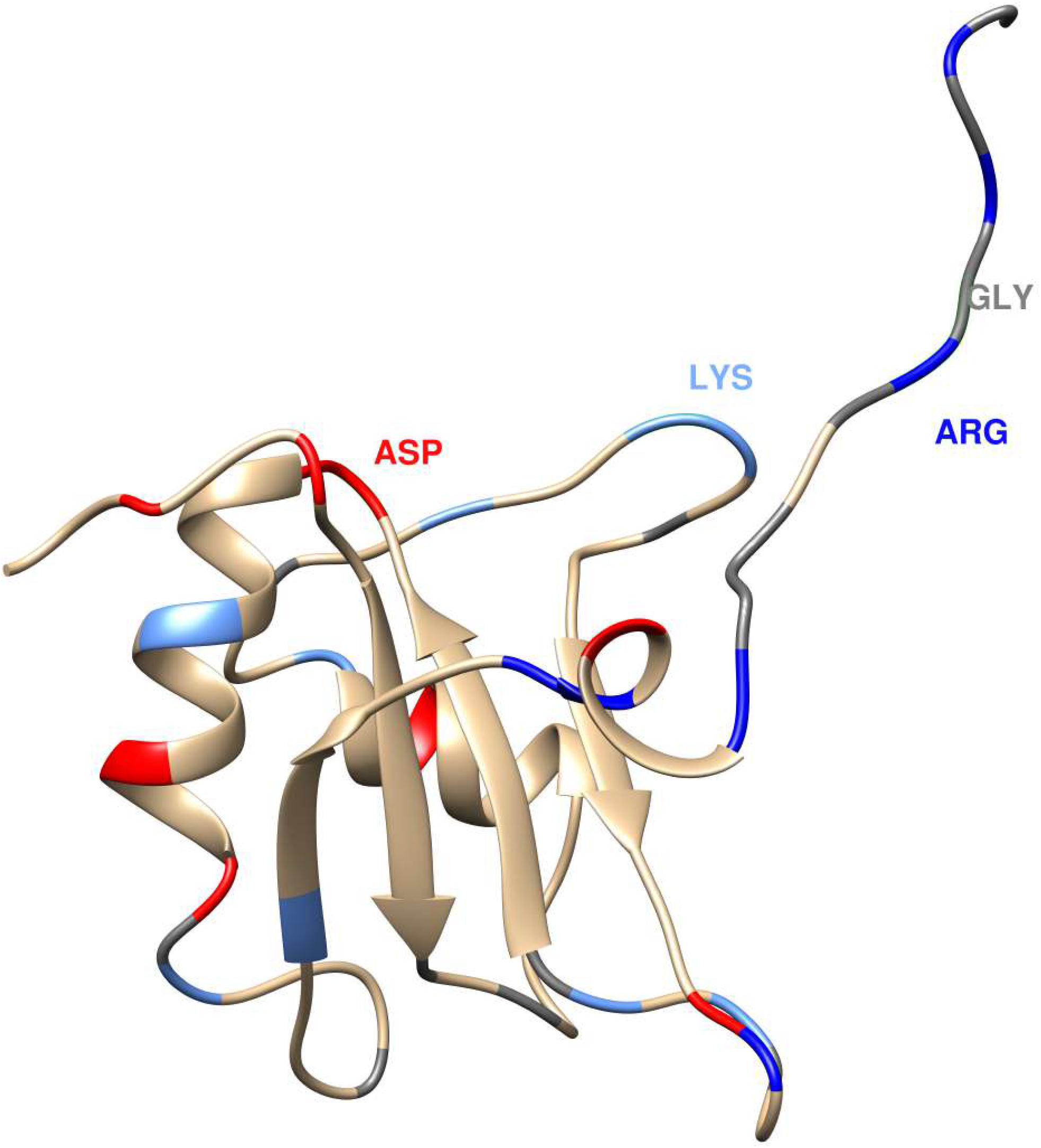
The location of key interacting residues Arg (blue), Lys (light blue), Asp (red), and Gly (gray) in the three-dimensional structure of RRM and RGG2 is depicted in FUS_390_ structure.

**FIG. S8:**
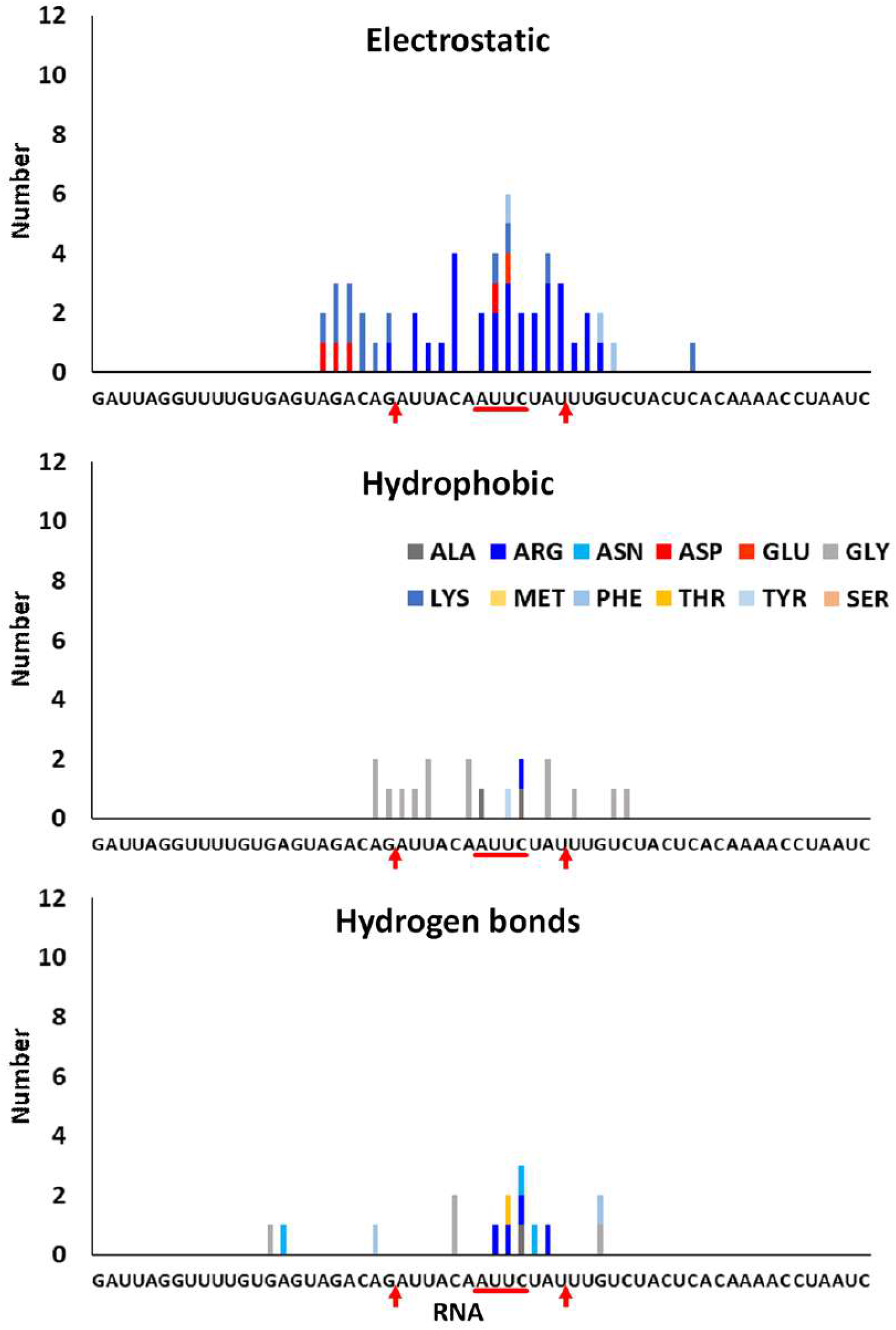
Amino acid-wise interaction histogram depicting the number of electrostatic, hydrophobic, and hydrogen bond interactions by each amino acid in *FUS*_418_. The number of interactions for each amino acid includes both RRM and RGG2

**FIG. S9:**
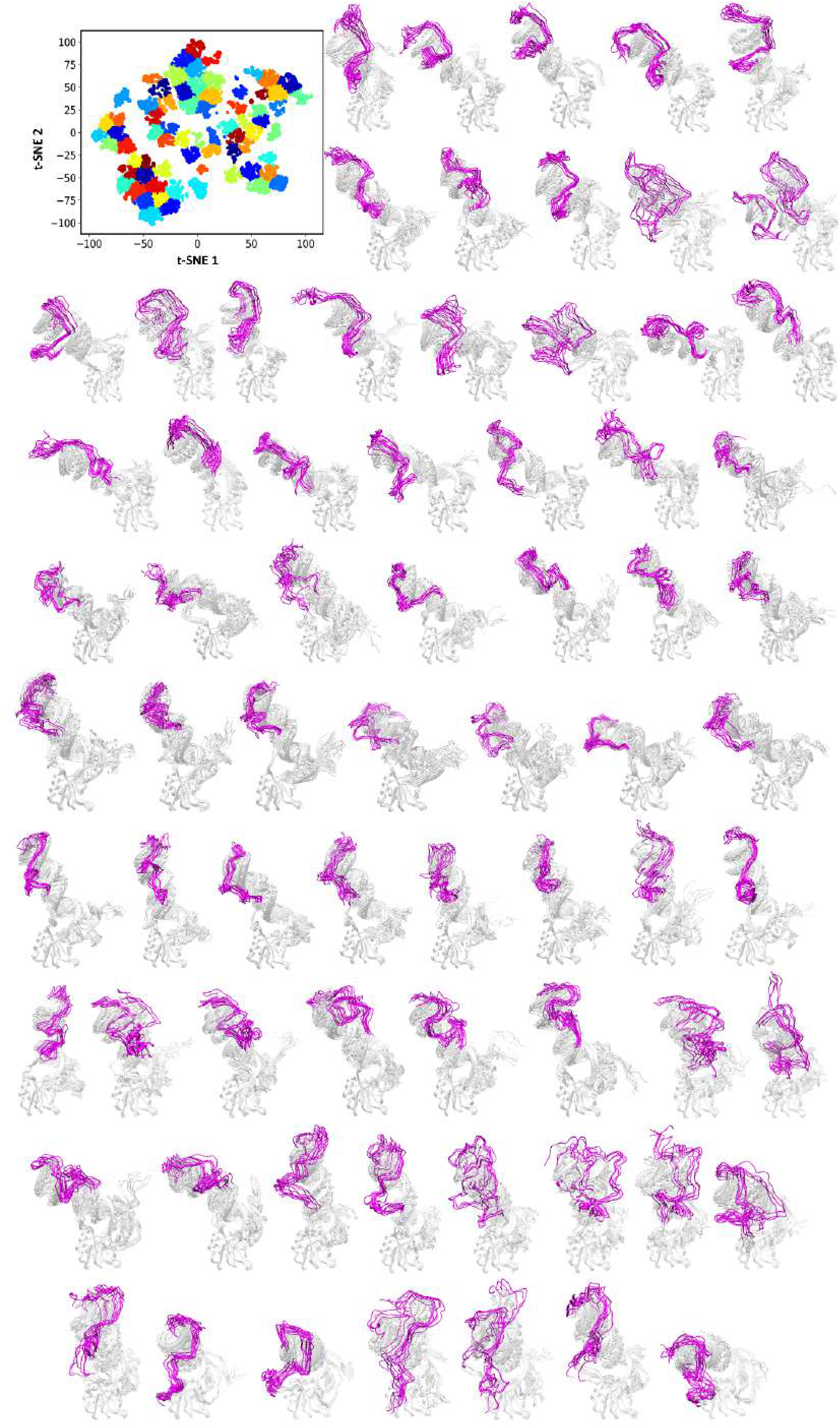
Clustering of the *FUS*_223*−*418_ ensemble by t-SNE and kMeans methods. The projection of the first two tSNE components classifies the sampled conformations into 70 distinct and unique clusters. 10 conformers from each cluster are superimposed and shown.

**TABLE S1.** Average lifetime of interactions calculated per residue with the AUUC (AAUG in case of RNAmut systems) motif in one of the three simulation trajectories of all the studied systems. The lifetimes were calculated by averaging the lifetimes of all-atom pairs per residue within a distance of 7 Å, normalized by the total number of contacts per residue-base pair.

| Systems | A | U/A | U | C/G |
| --- | --- | --- | --- | --- |
| NMR | Arg328 | Thr326, Arg328,<br>Lys334, Asn376 | Phe288, Lys334,<br>Thr370, Arg372 | Asp283, Asn285,<br>Arg371, Arg372,<br>Ala373 |
| $FUS_{RRM}$ | Arg328 (11%) | Thr326 (64%),<br>Arg328 (43%),<br>Lys334 (33%),<br>Arg372 (15%) | Phe288 (67%),<br>Lys334 (23%),<br>Thr370 (70%) | Asn323 (23%),<br>Tyr325 (30%) |
| $FUS_{380}$ | Arg328 (13%) | | | |
| $FUS_{385}$ | Arg372 (33%) | Arg328 (68%),<br>Lys334 (46%),<br>Arg372 (43%) | Phe288 (65%),<br>Tyr325 (54%),<br>Lys334 (21%),<br>Thr370 (68%),<br>Arg372 (44%),<br>Phe375 (28%) | Thr286 (63%),<br>Tyr325 (29%),<br>Thr338 (39%),<br>Ala369 (49%),<br>Arg371 (72%),<br>Arg372 (31%) |
| $FUS_{390}$ | Ala373 (33%),<br>Asn376 (21%),<br>Gly380 (6%) | Tyr325 (26%),<br>Arg328 (54%),<br>Arg372 (37%) | Phe288 (60%),<br>Thr370 (69%),<br>Arg372 (48%) | Thr286 (64%),<br>Thr338 (38%),<br>Arg371 (73%) |
| $FUS_{418}$ | Ala373 (72%),<br>Asp374 (31%) | Thr326 (33%),<br>Arg328 (61%),<br>Arg372 (50%) | Phe288 (44%),<br>Thr370 (67%),<br>Arg372 (58%) | Asn284 (35%),<br>Arg371 (75%),<br>Ala369 (40%) |
| $FUS_{223-418}$ | Ala373 (41%) | Arg328 (32%) | Phe288 (51%),<br>Asn323 (33%),<br>Tyr325 (64%),<br>Thr370 (73%) | Met321 (58%),<br>Arg371 (19%),<br>Arg372 (50%),<br>Ala373 (44%) |
| $FUS_{418} - RNA_{mut}$ | Arg386 (49%),<br>Arg372 (52%),<br>Ala373 (41%) | Arg328 (66%) | Phe288 (48%),<br>Asn323 (26%),<br>Tyr325 (44%),<br>Thr338 (55%),<br>Ala369 (66%),<br>Thr370 (69%) | Ala369 (53%),<br>Arg371 (81%),<br>Arg372 (41%) |
| $FUS_{390} - RNA_{mut}$ | Arg372 (43%) | Lys334 (25%),<br>Arg372 (38%) | Phe288 (67%),<br>Tyr325 (47%),<br>Lys334 (18%),<br>Thr370 (69%),<br>Arg372 (56%) | Asn284 (42%),<br>Ala369 (56%),<br>Thr370 (52%),<br>Arg371 (65%),<br>Arg372 (32%) |

## Notes

https://indianinstituteofscience-my.sharepoint.com/:f:/g/personal/bsangeetha_iisc_ac_in/Er0MtllWY-9EnxpAum3fs6MBDYJ_m3JHQzt7Bl4uIF-YyQ?e=RbG79d

### Competing Interest Statement

The authors have declared no competing interest.

### Summary of Updates

Polished up the draft.

